# Lidocaine Induces Apoptosis in Head and Neck Squamous Cell Carcinoma Through Activation of Bitter Taste Receptor T2R14

**DOI:** 10.1101/2022.11.22.517413

**Authors:** Zoey A. Miller, Arielle Mueller, Jennifer F Holivert, Ray Z. Ma, Sahil Muthuswami, April Park, Derek B. McMahon, Ryan M. Carey, Robert J. Lee

## Abstract

Head and neck squamous cell carcinomas (HNSCCs) have high mortality and significant treatment-related morbidity. It is vital to discover effective, minimally invasive therapies that improve survival and quality of life. Bitter taste receptors (T2Rs) are expressed in HNSCCs and their activation can induce apoptosis. Lidocaine is a local anesthetic used in various clinical applications which can also activate bitter taste receptor 14 (T2R14). Lidocaine may have some anti-tumor effects in other cancers, but the mechanism has been mysterious. Here, we found that lidocaine causes intracellular Ca^2+^ mobilization through activation of T2R14 in HSNCC cells. T2R14 activation with lidocaine depolarizes the mitochondrial membrane, inhibits cell proliferation, and induces apoptosis. Concomitant mitochondrial Ca^2+^ influx, ROS production causes T2R14-dependent accumulation of poly-ubiquinated proteins, suggesting inhibition of the proteasome as a novel component of T2R14-induced apoptosis. Lidocaine may have therapeutic potential in HNSCC as a topical gel or intratumor injection, warranting future clinical studies.

**HIGHLIGHTS:**

- Lidocaine activates bitter taste receptor 14 (T2R14) to increase intracellular Ca^2+^ and decrease cAMP in head and neck squamous cell carcinoma (HSNCC) cells
- Lidocaine decreases cell viability and cell proliferation, depolarizes the mitochondrial membrane, and causes production of reactive oxygen species (ROS)
- T2R14 activation with lidocaine induces apoptosis and inhibits the ubiquitin proteasome system (UPS)
- Lung squamous cell carcinoma (SCC) cells and HNSCC tumor spheroids undergo apoptosis or structural disbandment, respectively, with lidocaine

## INTRODUCTION

Head and neck squamous cell carcinomas (HNSCCs) arise in the mucosa of oral and nasal cavities, larynx, and pharynx, typically as a result of exposure to environmental carcinogens and/or the human papilloma virus.^1–4^ HNSCCs account for roughly 900,000 new cancer diagnoses per year, with a 5-year mortality rate of ∼50%.^1^ This is a result of frequent late-stage diagnosis, lack of preventive screening, and high rates of metastasis.^5–7^ Surgery, radiation, chemotherapy, and occasionally immunotherapy, are possible treatment options. Patients suffer from side effect burden, which negatively impacts their quality of life (QoL).^3^ Long-term survivors after effective treatment can be left with physical deformities, chronic pain, tracheostomy or feeding tube dependence, and inability to orally communicate, amongst other impacts.^8,9^ It is vital to develop new targeted therapies to effectively treat HNSCC while maintaining QoL.

A majority of HNSCC cases are localized to the oral cavity and oropharynx.^10^ A main function of this region is taste perception. Humans can perceive five tastes: sour, salty, sweet, umami, and bitter.^11^ Sweet, umami, and bitter flavors are perceived through the activation of specialized G protein-coupled receptors (GPCRs).^12–16^ GPCR taste receptors are classified into two groups: taste family 1 (T1Rs; sweet and umami) and taste family 2 receptors (T2Rs; bitter).^17^ While there are only 3 isoforms of T1Rs, there are 25 T2R isoforms in humans, which primarily function to protect against ingestion of bitter tasting toxins.^18^ The canonical T2R signaling pathway causes an intracellular calcium (Ca^2+^) increase and cyclic adenosine monophosphate (cAMP) decrease.^19^ This Ca^2+^ response is through activation of PLCβ2 via Gβγ subunits of the G-protein heterotrimer^20–22^, which produces inositol triphosphate (IP_3_) and stimulates the release of Ca^2+^ from the endoplasmic reticulum (ER).^23,24^

Beyond their role in bitter taste perception, T2Rs are involved in innate immunity, thyroid function, cardiac physiology, and other biologic processes.^25–29^ Furthermore, T2Rs have been studied in cancers including ovarian, breast, and HNSCC.^30–32^ Of the 25 T2R isoforms, T2R14 is being studied most actively.^33,34,35^ We previously showed that bitter agonists that activate various T2Rs cause an increase in nuclear and mitochondrial Ca^2+^ that leads to apoptosis in squamous de-differentiated non-ciliated airway epithelial cells and HNSCC cells.^21,32^ While we found that Ca^2+^ is required for this apoptosis, the underlying mechanisms of this T2R-induced endpoint are largely unknown despite other studies also linking T2Rs to apoptosis.

Lidocaine is commonly used as topical or injectable local for various clinical applications. ^36^ The compounds blocks sodium (Na^+^) channels to inhibit pain signals from sensory neurons.^37^ Some studies suggest that lidocaine may also have chemotherapeutic effects and decrease the rates of metastasis in other cancer types.^38–43^ However, no mechanism(s) have been identified for why lidocaine has these effects. Interestingly, lidocaine is bitter and interact with heterologously-expressed T2R14.^44^ From this, we hypothesized that that T2R14 and its agonist lidocaine may have pro-apoptotic effects in HNSCCs.

Here, we show that lidocaine induces Ca^2+^ responses in HNSCC cells through T2R14 activation. Moreover, lidocaine decreases HNSCC cell viability, depolarizes the mitochondrial membrane potential, elevates mitochondrial reactive oxygen species (ROS), and induces apoptosis. Apoptosis is blocked by T2R14 antagonism. Interestingly, we also observe that bitter agonists lidocaine can inhibit the proteasome, for the first time revealing a possible mechanism of T2R agonist-evoked apoptosis. Our data identify a new mechanism of action for the apoptotic effects of lidocaine in cancer cells and suggest that lidocaine may be useful to repurpose as a therapeutic for HNSCCs, either as a topical gel or intratumor injection.

## RESULTS

### Lidocaine stimulation causes an intracellular Ca^2+^ response

T2R14 is one of 25 T2R isoforms, and T2R14 is strongly expressed in oral and extraoral tissues.^45^ Lidocaine activation of T2R14 has only ever been recorded with heterologous expression. To understand if lidocaine can activate endogenous T2R14, Ca^2+^ responses were recorded in living HNSCC cells loaded with Fluo-4 AM.^46^ Dose-dependent Ca^2+^ responses were recorded in SCC47 oral cancer cells exposures to lidocaine at concentrations ranging from 0 to 10 mM, with 10 mM evoking the highest response (Figures 1A and 1B).^47,48^ Lidocaine-induced Ca^2+^ responses were compared to those with other bitter agonists. At both 5 and 10 mM, lidocaine caused higher Ca^2+^ responses than denatonium benzoate (Figures 1C and 1D). Denatonium benzoate is a structurally-similar non-T2R14 bitter agonist that also can induce apoptosis in HNSCC cells; denatonium benzoate is considered one of the most bitter compounds due to its activation of eight T2R isoforms (T2R4, 8, 10, 13, 39, 43, 46, and 47 agonist).^49,50^ Lidocaine also induced the highest Ca^2+^ response among other T2R agonists screened at their respective maximum concentrations in aqueous solution, including denatonium benzoate, thujone (T2R14 agonist), flufenamic acid (T2R14 agonist), and ATP (purinergic receptor agonist) in SCC47, FaDu pharyngeal cancer, and RPMI2650 nasal squamous carcinoma cells(Figures 1E-1G).^34,44,51^ Lidocaine also induced Ca^2+^ responses at 10 mM in SCC4 and SCC90 oral cancer cells that were comparable to FaDu and SCC47 cells (Figure 1H-1K). Unlike lidocaine, closely related anesthetic procaine, not known to activate T2R14^52^, did not induce Ca2+ responses when tested at equivalent concentrations (Figure 1K). Thus, the response to lidocaine is not due to Na^+^ channel inhibition but likely from intracellular Ca^2+^ sources.

**Figure 1.**
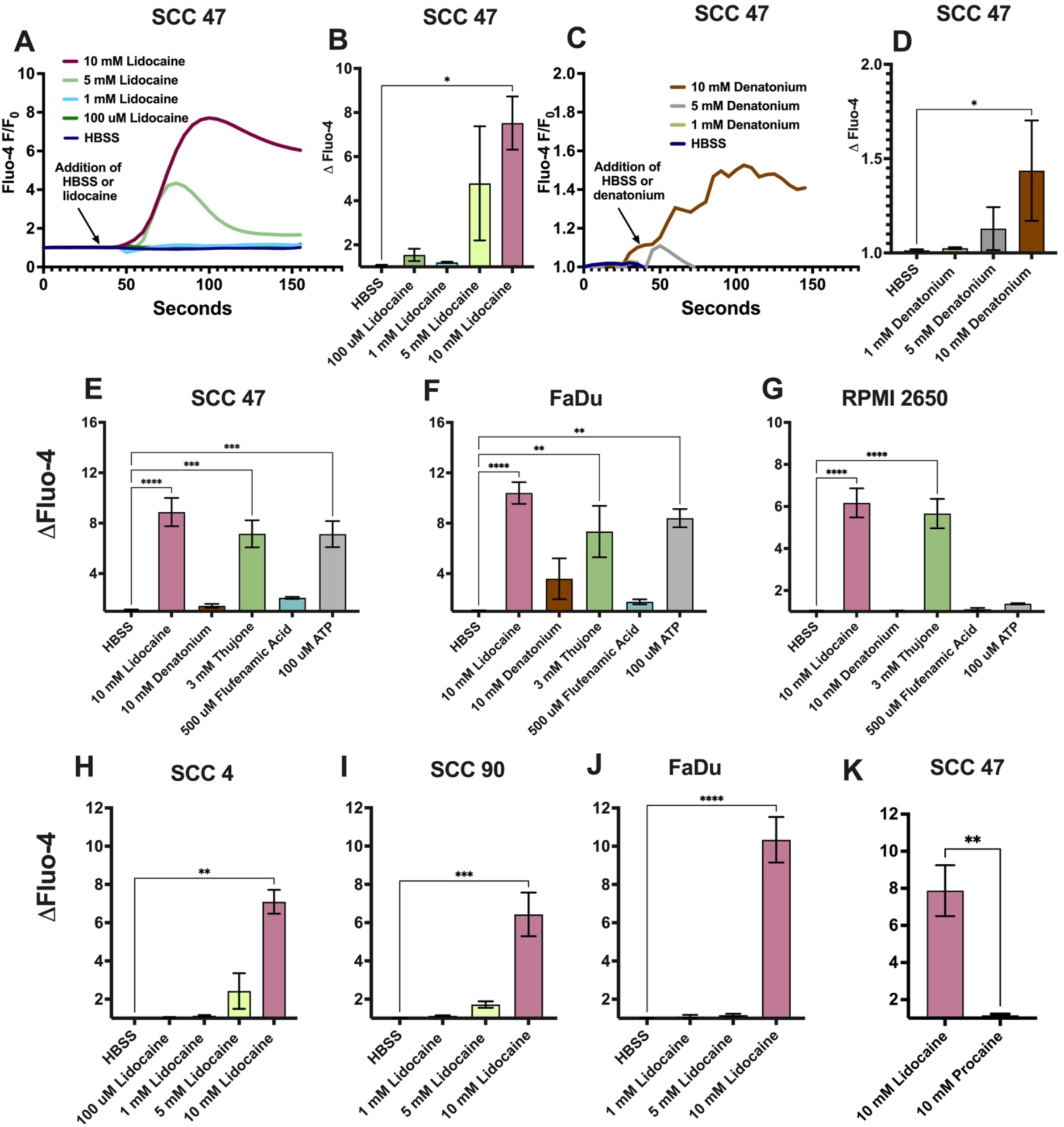
Lidocaine induces Ca^2+^ responses in HNSCC cells. HNSSC cell lines SCC 47, FaDu, RPMI 2650, SCC 4, and SCC 90 were loaded with Ca^2+^ indicator Fluo-4 and imaged for subsequent Ca^2+^ responses with bitter agonists **A-B)** Representative SCC 47 Ca^2+^ response over time (*A*) and peak Ca^2+^ responses with 0 – 10 mM lidocaine; mean ± SEM with >3 experiments using separate cultures (*B*). Significance by 1-way ANOVA with Bonferroni posttest comparing HBSS to each concentration of lidocaine. **C-D)** Representative SCC 47 Ca^2+^ response (*C*) over time and peak fluorescent Ca^2+^ responses with 0 – 10 mM denatonium benzoate (*D*). Peak fluorescence mean ± SEM with >3 experiments using separate cultures. Significance by 1-way ANOVA with Bonferroni posttest comparing HBSS to each concentration of denatonium. **E-G)** SCC 47 (*E*), FaDu (*F*), and RPMI 2650 (*G*) peak Ca^2+^ responses with HBSS, 10 mM lidocaine, 10 mM denatonium benzoate, 15 mM denatonium benzoate, 3 mM thujone, 500 µM flufenamic acid, or 100 µM ATP. Mean ± SEM with >3 experiments using separate cultures. Significance by 1-way ANOVA with Bonferroni posttest comparing HBSS to each agonist **H-J)** SCC 4 (*H*), SCC 90 (*I*), and FaDu (*J*) peak fluorescent Ca^2+^ responses with 0 – 10 mM lidocaine; mean ± SEM with >3 experiments using separate cultures. Significance by 1-way ANOVA with Bonferroni posttest comparing HBSS to each concentration of lidocaine **K)** SCC 47 peak Ca^2+^ response with 10 mM lidocaine or procaine; mean +/- SEM with >3 experiments using separate cultures. Significance by unpaired t-test. P < 0.05 (*), P < 0.01 (**), P < 0.001 (***), and no statistical significance (ns).

Stimulation with lidocaine dose-dependently reduced Ca^2+^ responses with secondary ATP stimulation. ATP activates intracellular Ca^2+^ responses via purinergic GPCRs.^51^ ATP (100 µM) was used following lidocaine (1 – 10 mM) stimulation in SCC 4 and SCC 47 cells. As lidocaine concentration increased, the post-ATP Ca^2+^ response decreased, suggesting depletion of Ca^2+^ stores by lidocaine (Figures S1A-F).

We further tested if the lidocaine Ca^2+^ response was from mobilization of intracellular Ca^2+^ stores or extracellular Ca^2+^ influx from plasma membrane Ca^2+^ channels. Ca^2+^-linked GPCRs, including T2Rs, most often induce intracellular Ca^2+^ responses primarily originating from the ER inositol trisphosphate (IP_3_) receptor (IP_3_R) Ca^2+^ release, with a sustained Ca^2+^ responses requiring store-operated Ca^2+^ entry.^53–56^ We recorded Fluo-4 responses in HBSS with or without extracellular Ca^2+^. The Ca^2+^-free buffer contained no added Ca^2+^ plus EGTA to chelate any residual free extracellular Ca^2+^. Cells stimulated with lidocaine in HBSS without Ca^2+^ had a similar initial Ca^2+^ peak as cells stimulated in the presence of extracellular Ca^2+^ (Figure 2A). However, there was a faster rate of returning to baseline fluorescence compared with cells stimulated in HBSS with Ca^2+^ (Figure 2A). This shows that the initial Ca^2+^ response observed with lidocaine is intracellular release (Figure 2B). In addition, kb-r7943, a Na^+^/Ca^2+^ membrane exchanger inhibitor, had no effect on lidocaine-induced Ca^2+^ response, further indicating mobilization from intracellular stores rather than Ca^2+^ influx (Figure S2A).

**Figure 2.**
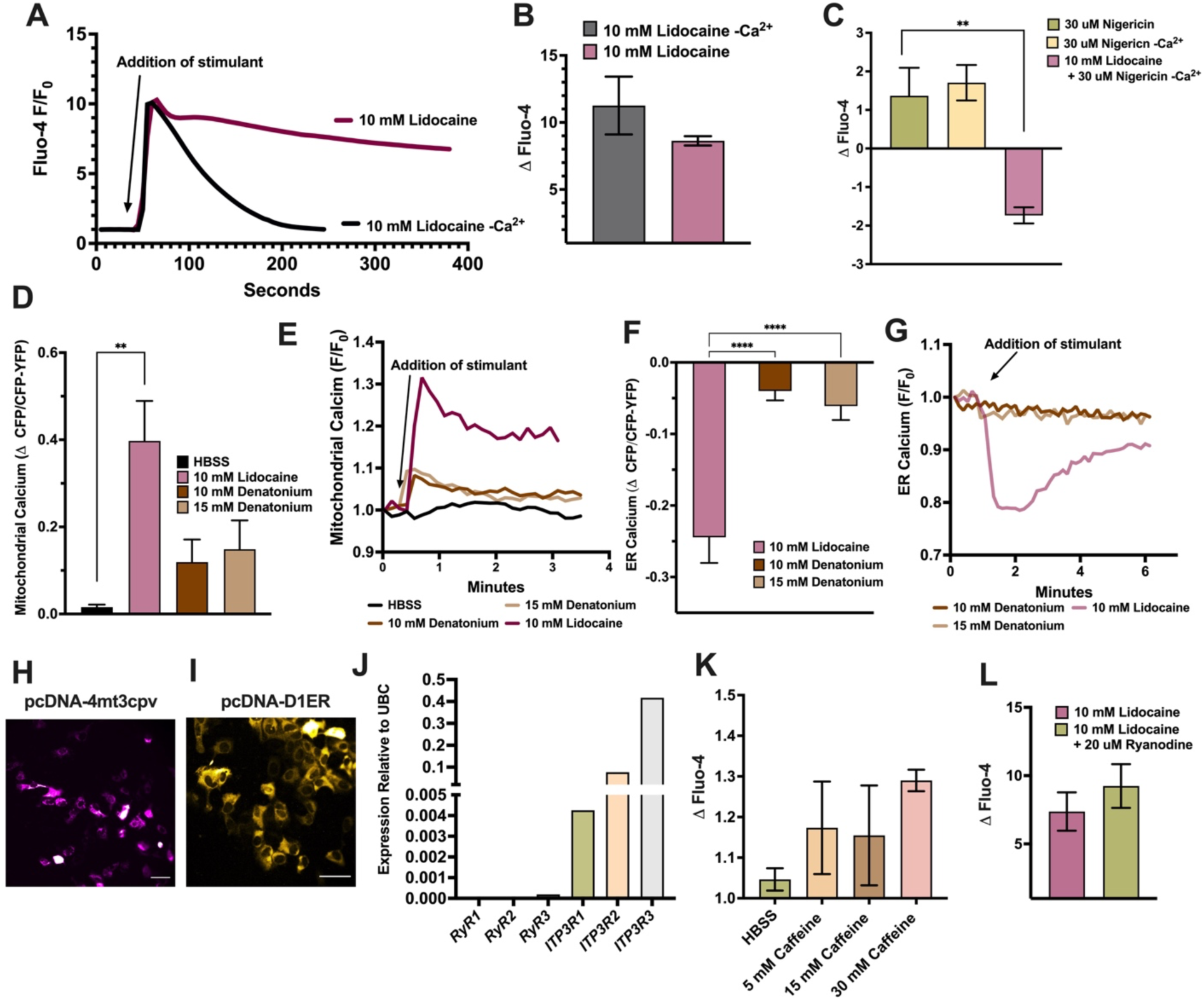
Ca^2+^ response with lidocaine is due to intracellular Ca^2+^ release. SCC 47 cells were loaded with Fluo-4 and imaged for subsequent Ca^2+^ responses with bitter agonists. **A-B)** Representative SCC 47 Ca^2+^ trace (*A*) and peak Ca^2+^ responses (*B*) with 10 mM lidocaine in HBSS ± extracellular Ca^2+^. Peak fluorescence mean +/- SEM with >3 experiments using separate cultures. No significant difference by t-test **C)** SCC 47 peak Ca^2+^ response with nigericin stimulation in the presence or absence of extracellular Ca^2+^ and following stimulation with 10 mM lidocaine in the absence of extracellular Ca^2+^. Significance by 1-way ANOVA with Bonferroni posttest comparing nigericin response to nigericin responses in -Ca^2+^ and post-lidocaine stimulation. **D-G)** SCC 47 cells were transfected with pcDNA-4mt3cpv (*D*-*E*) or pcDNA-D1ER (*F*-*G*) 24 – 48 hours prior to Ca^2+^ imaging. **D-E)** SCC 47 mitochondrial (pcDNA-4mt3cpv) peak (mean ± SEM; *D*) and representative trace (E) CFP/CFP-YFP fluorescent Ca^2+^ responses with HBSS, 10 mM lidocaine, 10 mM denatonium benzoate, or 15 mM denatonium benzoate; >3 experiments using independent cultures. Significance by 1-way ANOVA with Bonferroni posttest comparing HBSS to each agonist. **F-G)** SCC 47 ER (pcDNA-D1ER) change in fluorescence (*F*) and trace (*G*) CFP/CFP-YFP fluorescent Ca^2+^ responses with 10 mM lidocaine, 10 mM denatonium benzoate, or 15 mM denatonium benzoate. Peak fluorescence mean ± SEM with >3 experiments using separate cultures. Significance by 1-way ANOVA with Bonferroni posttest comparing lidocaine to each concentration of denatonium. **H-I)** SCC 47 representative image of pcDNA-4mt3cpv (*H*) and pcDNA-D1ER CFP/YFP (*I*) baseline localization. Scale bar = 30 µm. **J)** mRNA expression of ER Ca^2+^ channels in SCC 47 relative to UBC. **K)** SCC 47 peak Ca^2+^ responses with 0-30 mM RyR agonist caffeine; mean ± SEM with >3 experiments using separate cultures. Significance by 1-way ANOVA with Bonferroni posttest comparing HBSS to each concentration of caffeine. **L)** SCC 47 peak lidocaine Ca^2+^ responses ± prior ryanodine inhibition (1 hour). Fluorescent means ± SEM with >3 separate cultures. No significant difference by unpaired t-test. P < 0.05 (*), P < 0.01 (**), P < 0.001 (***), and no statistical significance (ns or no indication).

To begin to dissect out the origin of the Ca^2+^ release, cells were also stimulated with nigericin, which causes release of intracellular acidic Ca^2+^ stores, including lysosomal stores.^57^ Nigericin alone induced small Ca^2+^ responses, as expected. However, after lidocaine stimulation in the absence of extracellular Ca^2+^, nigericin no longer induced this response. This is most likely due to lidocaine depleting intracellular acidic Ca^2+^ stores (Figure 2C). However, the smaller magnitude of the nigericin-induced Ca^2+^ response vs the lidocaine-induced response suggested this was not the majority of lidocaine-induced Ca^2+^.

GPCR activation can also alter mitochondrial Ca^2+^ content.^58,59^ We used a mitochondrial-localized fluorescent Ca^2+^ biosensor, pcDNA-4mt3cpv, to measure mitochondrial Ca^2+^ dynamics^60^ (Figure 2D). Upon stimulation, a significant influx of Ca^2+^ into the mitochondria was observed with lidocaine (Figures 2D and 2E; 2H). The influx of Ca^2+^ into the mitochondria with lidocaine stimulation was significantly greater than with denatonium. An ER-specific fluorescent Ca^2+^ biosensor, pcDNA-D1ER, was used to measure ER Ca^2+^ dynamics (Figure 2F).^60^ In contrast to the mitochondria, lidocaine stimulation caused a significant decrease in ER Ca^2+^, also much larger compared to denatonium (Figures 2F and 2G; 2I).

It is likely that this is ER Ca^2+^ release. We measured the mRNA expression of ryanodine receptor (*RyR*) and inositol triphosphate receptor (IP_3_R) isoforms in SCC 47 cells. Cells had much higher expression of *ITP3R* compared to *RyR* (Figure 2J). There was also no Ca^2+^ response with caffeine, an agonist of *RyR*, and no change in Ca^2+^ responses with lidocaine in the presence of ryanodine, an antagonist of *RyR* (Figures 2K and 2L).^61,62^ Combined with the low expression of *RyR*, this indicates that Ca^2+^ release from the ER is most likely through IP_3_R.

### Lidocaine activates T2R14

Following GPCR activation, GTP-bound guanine nucleotide-binding proteins (G proteins) disassociate from their receptors and carry out effector actions.^63,64^ As GPCRs, T2Rs can couple to Gα_gustducin_ or Gα_i_, Gβ_1/3_ and Gγ.^15,23,65^ To test if lidocaine causes GPCR activation, cells were incubated with suramin, a pan-inhibitor of G protein association to their receptors.^66^ Inhibition of G proteins with suramin significantly dampened Ca^2+^ responses with lidocaine (Figure 3A). Cells were also incubated with pertussis toxin, an inhibitor of Gα_i/o_ and Gα_gustducin_ subunits.^67^ Cells treated with pertussis toxin had significantly dampened Ca^2+^ responses with lidocaine (Figure 3B). This suggests GPCR activation caused by lidocaine in HNSCC cells.

**Figure 3.**
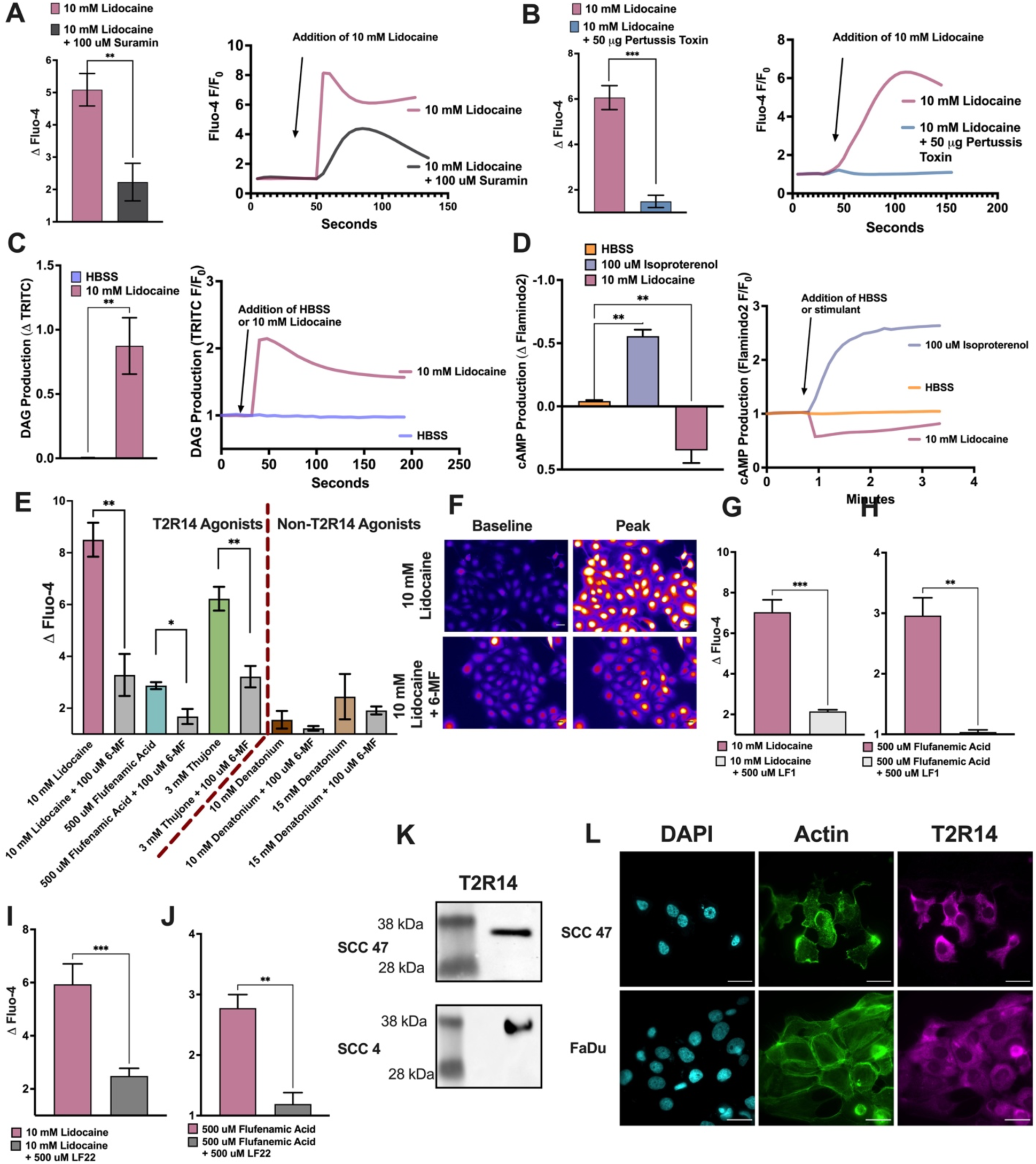
Lidocaine increases intracellular Ca^2+^ through activation of T2R14. **A)** SCC 47 peak Ca^2+^ (left; mean ± SEM) and representative trace with 10 mM lidocaine +/- 1-hour prior incubation with 100 µM suramin; n = >3 experiments using separate cultures. Significance by t-test. **B)** SCC 47 peak Ca^2+^ (left; mean ± SEM) and representative trace of 10 mM lidocaine +/- 18-hour prior incubation with 50 μg/ml pertussis toxin or media only; n = >3 experiments using separate cultures. Significance by t-test. **C)** SCC 47 cells were transduced with fluorescent DAG biosensor and imaged for subsequent DAG responses with 10 mM lidocaine. SCC 47 peak DAG (left; mean ± SEM) and representative DAG trace (right) with HBSS or 10 mM lidocaine; n = >3 experiments using independent cultures. Significance by t-test. **D)** SCC 47 cells were transfected with cAMP biosensor Flamindo2 and imaged for subsequent cAMP responses with HBSS, 10 mM lidocaine or 100 µM isoproterenol. SCC 47 peak cAMP response (left; mean ± SEM) and representative cAMP traces (right) with HBSS, 100 µM isoproterenol, or 10 mM lidocaine; n = >3 experiments using separate cultures. Significance by 1-way ANOVA with Bonferroni’s posttest comparing each agonist to HBSS. **E)** SCC 47 peak Ca^2+^ responses with bitter agonists ± prior 1-hour incubation with 100 µM 6-methoxflavanone (6-MF); mean +/- SEM with >3 experiments using separate cultures. Significance by 1-way ANOVA with Bonferroni’s posttest comparing each agonist ±6-MF. **F)** Representative images of SCC 47 peak Ca^2+^ responses with lidocaine ± 6-MF. Scale bar = 30 µm. **G-H)** SCC 47 peak Ca^2+^ responses with 10 mM lidocaine (*G*) or 500 uM flufenamic acid (*H*) ± prior 1-hour incubation with 500 uM LF1. **I-J)** SCC 47 peak Ca^2+^ fluorescent responses with 10 mM lidocaine (*I*) or 500 uM flufenamic acid (*J*) ± prior 1-hour incubation with 500 uM LF22; mean +/- SEM with >3 separate independent cultures. Significance by unpaired t-test. T2R14 primary antibody (1:500 or 1:1000) was used to detect expression in SCC 47, SCC4 and FaDu cells. **K)** T2R14 protein expression in SCC 47 and SCC 4 cells via Western blot (1:500 primary antibody). **L)** T2R14 expression in SCC 47 and FaDu cells via immunofluorescence stain with DAPI (nucleus) and phalloidin (actin) (1:100 primary antibody). P < 0.05 (*), P < 0.01 (**), P < 0.001 (***), and no statistical significance (ns or no indication).

Interestingly, cells incubated with pertussis toxin and stimulated with denatonium had no significant differences in Ca^2+^ responses (Figure S2B). Cells were also incubated with YM254890, a Gαq/11 selective inhibitor.^68^ While YM25480 decreased Ca^2+^ responses with denatonium, it did not affect responses with lidocaine (Figure S2C). shRNA targeting T2R4 significantly downregulated Ca^2+^ responses with denatonium, while leaving responses with lidocaine unchanged (Figures S2D and S2E). These data support previous studies that lidocaine interacts with a separate T2R isoforms than denatonium.^44^

Canonical GPCR activation of phospholipase C (PLC) also produces diacylglycerol (DAG).^69^ To test if lidocaine stimulates DAG production, an RFP DAG biosensor was transduced via BacMam into cells. Lidocaine stimulation significantly increased DAG production compared to HBSS alone (Figure 3C). In addition, canonical T2R Gαi-coupling would be expected to be accompanied by a decrease in cAMP, which was measured using a fluorescent cytosolic cAMP biosensor, Flamindo2.^70^ Lidocaine significantly decreased cAMP compared to HBSS alone (Figure 3D). Isoproterenol, a β-adrenergic agonist, was used as a positive control to increase cAMP. The observed PTX-sensitive, suramin-sensitive lidocaine intracellular Ca^2+^ responses coincide with the timing of both increases in DAG and decreases in cAMP, all together confirming that lidocaine activates a Gα_i/o/gustudin_-coupled GPCR.

To understand if lidocaine interacts with T2R14 specifically, cells were incubated with 100 µM 6-methoxyflavonone (6-MF), a T2R14 antagonist.^71^ Incubation with 6-MF significantly decreased Ca^2+^ responses with lidocaine (Figure 3E and 3F). 6-MF also significantly dampened responses with flufenamic acid and thujone, two other T2R14 agonists (Figure 3E). 6-MF did not alter Ca^2+^ with denatonium, a non-T2R14 agonist (Figure 3E). A recent preprint also identified several other structurally-distinct T2R14 inhibitors.^72^ We tested two of these inhibitors, LF1 and LF22, and found that they also blocked the Ca^2+^ responses to both lidocaine and flufenamic acid (Figures 3G-3J). LF-1 and LF22 also reduced Ca^2+^ responses to other T2R14 agonists (but not to denatonium or ATP-non T2R14 agonists) in H292 cells, previously shown to express T2R14, and in FaDu cells (Figure S3) ^73^. Ca^2+^ responses to lidocaine were also dampened when T2R14 expression was decreased in HEK 293T, which endogenously express T2R14, cells using shRNA (Figure S4) ^21^.

Several HNSCC cell lines express T2R14 by qPCR.^32^ To further validate expression of T2R14, we measured protein levels via Western blot, immunofluorescence, and mRNA in several HNSCC cell lines used here (Figures 3K and 3L; Figure S5A and S5B). T2R14 was present in SCC 47, FaDu, and SCC 4 cells. These data indicate that lidocaine interacts with a GPCR in HNSCC cells, likely T2R14, to trigger an intracellular Ca^2+^ response.

### Lidocaine decreases cell viability and mitochondrial membrane potential

Lidocaine is being investigated as a therapy for several cancers, but its mechanism of action is unknown.^43,48,74^ After establishing an interaction between T2R14 and lidocaine, we tested if lidocaine had anti-proliferative effects in HNSCC cells. Using a crystal violet assay, we found that 10 mM lidocaine significantly decreased cellular proliferation after just 6 hours (Figure S6A). After 24 hours, concentrations as low as 1 mM significantly decreased cellular proliferation (Figure S6B).

To measure cell viability, we used an XTT assay as an indirect measurement of cellular NADH production.^75^ Lidocaine (5 mM and 10 mM) and denatonium reduced cell viability in SCC 47 cells (Figure 4A). This was observed after only 2 hours, with the full assay covering 6 hours. These results were also observed in both FaDu and SCC 4 cells. FaDu cells were more sensitive to denatonium benzoate than lidocaine (Figure 4B). Conversely, SCC 4 cells were more sensitive to lidocaine than denatonium (Figure S6C). These data suggest that different types of HNSCCs may have different sensitivities to specific T2R agonists. However, lidocaine was effective at reducing XTT absorbance in all cell types tested. TAKE OUT 6-MF XTT

**Figure 4.**
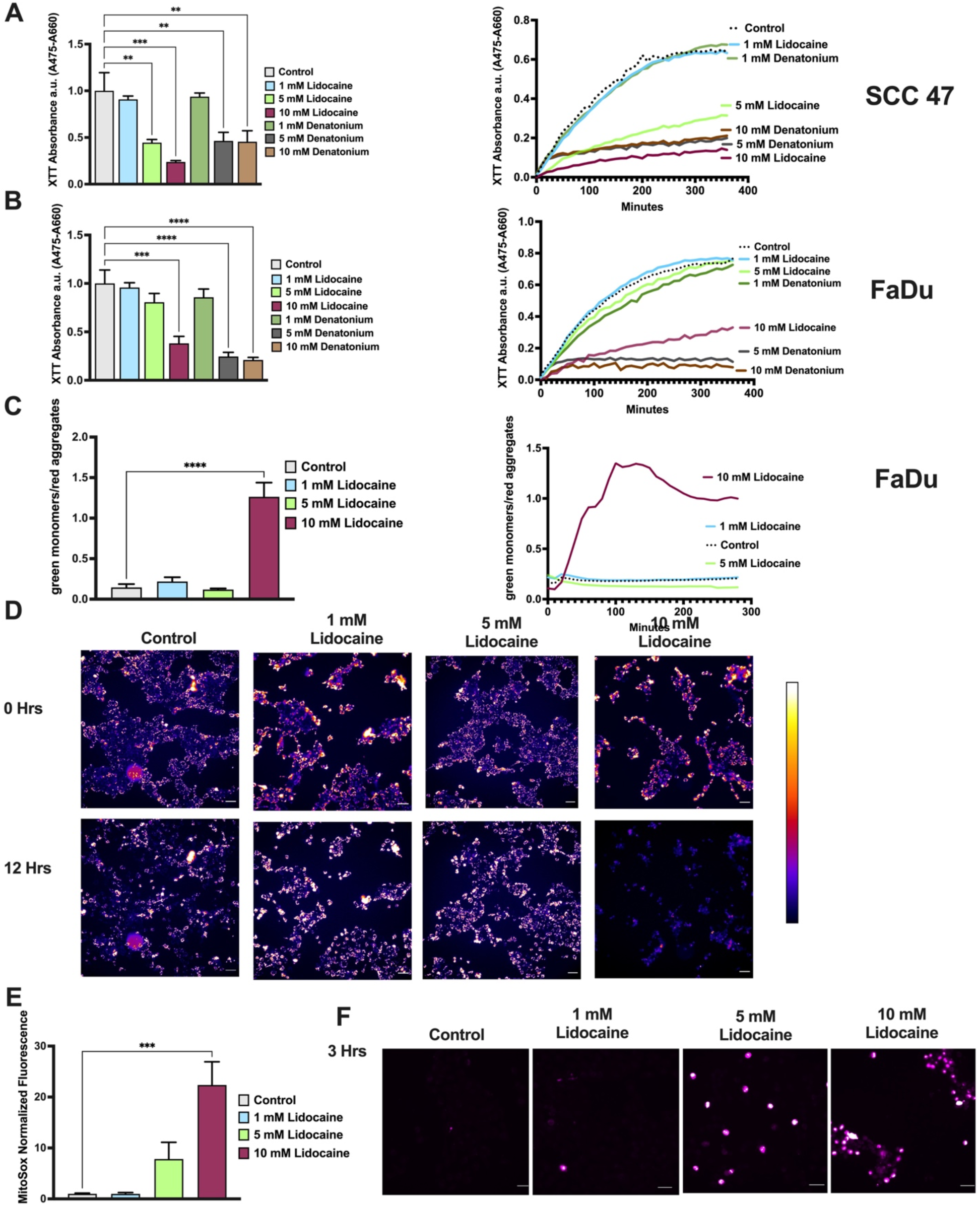
Lidocaine decreases cell viability and depolarizes the mitochondrial membrane. SCC 47 and FaDu cells were incubated with bitter agonists with XTT dye, an indicator of NADH production. A decrease in the difference of absorbance (475 nm – 660 nm) indicates reduced NADH production. **A)** SCC 47 and **B)** FaDu XTT absorbance values (475 nm – 660 nm) at 120 minutes of incubation with 0 – 10 mM lidocaine, or 0 -10 denatonium. Absorbance values were measured over six hours as seen traces (n=1). SCC 47 cells were loaded with fluorescent JC-1 dye 15 minutes prior to incubation with bitter agonists. An increase in FITC (dye monomers) / TRITC (dye aggregates) represents depolarized mitochondrial membrane potential. **C)** FaDu fluorescent green/red (FITC/TRITC channel emission ratio) ratio at 100 minutes of incubation with 0 – 10 mM lidocaine (mean ± SEM) and representative trace; n = >3 separate cultures. Significance by 1-way ANOVA with Bonferroni posttest comparing media to each concentration of lidocaine. **D)** FaDu representative images of change in red dye aggregates (indicative of loss of mitochondrial membrane potential) at 0 and 12 hours with 0 – 10 mM lidocaine. SCC 47 cells were treated with 0-10 mM lidocaine for 3 hours and then loaded with MitoSox superoxide indicator dye. **E)** SCC 47 MitoSox fluorescent values at 3 hours post stimulation. Fluorescent mean +/- SEM with >3 separate cultures. Significance by 1-way ANOVA with Bonferroni posttest comparing media to each concentration of lidocaine **F)** SCC 47 representative images of MitoSox fluorescence post-lidocaine. Scale bars = 30 µm. P < 0.05 (*), P < 0.01 (**), P < 0.001 (***), and no statistical significance (ns or no indication).

Despite Ca^2+^ mobilization observed with ATP in previous experiments (Figures 1E-1G), we wanted to test if ATP would also decrease cell viability. Concentrations ranging from 1 – 100 uM (saturating) ATP did not affect cell viability (Figure S6D), suggesting the T2R-activated Ca^2+^ responses are uniquely localized or paired with parallel signaling pathways that reduce viability.

Mitochondrial membrane potential (MMP) is an indicator of overall mitochondrial health, as it affects the bulk supply of ATP production.^76,77^ A hyperpolarized MMP reflects a healthy mitochondria, while the MMP will become depolarized in response to cellular stressors.^78,79^ We used a ratiometric JC-1 assay to assess mitochondrial health in HNSCC cells in response to lidocaine. This was measured using a ratio of JC1 monomers (green fluorescent) to aggregates (red fluorescent). Red aggregates accumulate in hyperpolarized mitochondria, and increased red-to-green ratio reflects hyperpolarization. We found that 10 mM lidocaine significantly depolarized the mitochondrial membrane (Figures 4C and 4D). These changes were observed as early as 100 minutes during the assay. Using MitoSox, a fluorescent mitochondrial superoxide indicator, we found that 10 mM lidocaine also produces reactive oxygen species (ROS) (Figures 4F and 4G). This highlights the potential anticancer effects of lidocaine.

### Lidocaine induces apoptosis via T2R14

Lidocaine induces apoptosis in a variety of cancerous cell types, however the mechanism has not yet been elucidated.^25,42,80–87^ We utilized a CellEvent assay to measure caspase-3 and -7 cleavage in HNSCC cells after exposure to lidocaine. The green CellEvent dye fluoresces upon cleavage with either caspase-3 or -7 if/when apoptosis occurs.^88–90^ This assay revealed that lidocaine induces apoptosis in SCC 47 and FaDu cells (Figure 5A and 5B; Figure S4A). SCC 47 cells were more sensitive to lidocaine-induced apoptosis than FaDu cells, as it took FaDu cells longer to undergo caspase cleavage (Figure 5B; Figure S7A). Nonetheless, lidocaine activated apoptosis in both cell types. Differential Interference Contrast (DIC) images were taken in parallel with the CellEvent assay in SCC 47 cells, revealing poor cellular morphology (rounding and lifting) coinciding with the apoptosis (Figure 5G).

**Figure 5.**
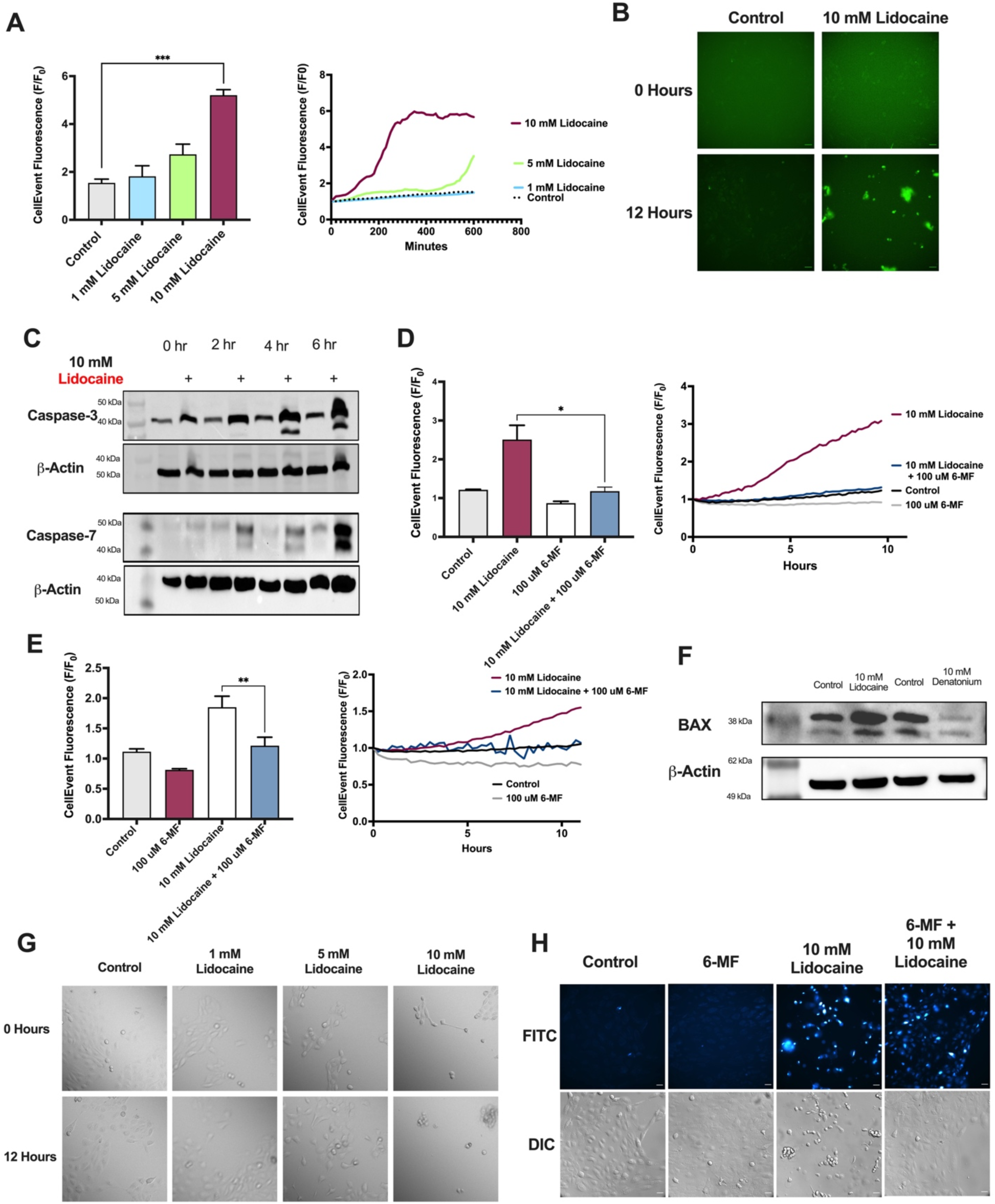
Lidocaine induces apoptosis via T2R14. **A)** SCC 47 CellEvent (caspase 3 and 7 indicator) fluorescence at 4 hours (mean ± SEM) with 0 - 10 mM lidocaine (left) and representative trace of CellEvent fluorescence over 12 hours with 10 mM lidocaine; n = >3 separate cultures on independent days. Significance determined by one-way ANOVA with Bonferroni posttest between control and lidocaine-treated cells. **B)** SCC 47 representative images of CellEvent caspase cleavage at 0 and 12 hours with 10 mM lidocaine. **C)** SCC 47 cells were treated with 10 mM lidocaine for 0, 2, 4, or 6 hours. Caspase-3, caspase-7, and β-Actin primary antibody (1:1000) were used with goat anti-rabbit or anti-mouse IgG-horseradish peroxidase secondary antibodies (1:1000). Immunoblots representative of 3 independent experiments using cells at different passage number on different days. **D-E)** SCC 47 (*D*) or FaDu (*E*) CellEvent fluorescence (peak mean ± SEM on left and representative trace on right) at 10 hours with 10 mM lidocaine ± 100 µM 6-MF. Significance was determined by unpaired t*-*test **F)** SCC 47 cells were treated with 10 mM lidocaine for 6 hours. Bax and β-Actin primary antibody (1:1000) were used with goat anti-rabbit or anti-mouse IgG-horseradish peroxidase secondary antibodies (1:1000). Immunoblots representative of 3 independent experiments using cells at different passage number on different days. **G)** Representative SCC 47 images of DIC cell morphology with 0-10 mM lidocaine. Scale bars = 30 uM. Representative of 3 independent experiments using cells at different passage number on different days. **F)** Representative DIC and Cell Event (green FITC fluorescence channel) images in SCC 47 cells with 0-10 mM lidocaine ± 100 uM 6-MF. Scale bars = 30 uM. Representative of 3 independent experiments using cells at different passage number on different days. P < 0.05 (*), P < 0.01 (**), P < 0.001 (***), and no statistical significance (ns or no indication).

To further verify caspase cleavage, full length and cleaved forms of both caspases were detected by Western blot in SCC 47 cells using 10 mM lidocaine over 6 hours. Caspase-7 cleavage occurred as early as 2 hours, while caspase-3 cleavage occurred as early as 4 hours (Figure 5C). In addition, Bcl-2-associated X (BAX), a pro-apoptotic protein, was activated with lidocaine stimulation, but not denatonium after 6 hours(Figure 5F).^91^ The two bitter agonists lidocaine and denatonium may have different underlying mechanisms of cell death induction, or activation of BAX with denatonium may be slower. Although BAX was activated with lidocaine, the mRNA levels of *BAX*, Bcl-2 agonist of cell death (*BAD*), and *Bcl-2* were unchanged (Figure S7B-D).

We tested if lidocaine could be translated into other cancers. We found that 10 mM lidocaine caused a Ca^2+^ response in NCI H520, a lung squamous cell cancer (SCC)-derived cell line, and induced apoptosis as well (Figure S7E and S7F). To better understand if the observed apoptosis was a direct function of T2R14 activation via lidocaine, we used the CellEvent assay with or without 6-MF (T2R14 antagonist). After 10 hours, 6-MF blocked apoptosis in SCC 47 and FaDu cells co-incubated with 10 mM lidocaine (Figures 5D and 5E). Complimentary to our original CellEvent data, DIC images revealed that 6-MF preserved overall cell morphology, even in the presence of lidocaine (Figure 5H). Lidocaine can induce apoptosis in HNSCCs via activation of a T2R, specifically T2R14, which could be potentially advantageous in treating HNSCCs or other cancers. Results from lung SCC broadens the scope of lidocaine’s potential use beyond HNSCC.

### Lidocaine may inhibit the proteasome as a mechanism of apoptosis

The exact mechanism by which lidocaine induces apoptosis in cancer cells is still unclear.^92–95^ In HNSCC at least, this depends on T2R14. We continued to studying the mechanisms of T2R14-induced apoptosis. When we Western blotted for caspase-3 and -7 cleavage as markers of apoptosis activation, we noticed there were increased cleaved and un-cleaved forms of the protein (Figure 5C). We tested if HNSCC cells were upregulating caspase- 3 and -7 production (i.e., mRNA transcription or protein translation) in response to lidocaine, or if there was inhibition of normal proteolytic degradation processes. There were no differences between caspase-3 or -7 (*CASP3 and CASP7)* mRNA at 0- and 6-hours post-lidocaine stimulation (Figure 6A). These proteins were then blotted for with 10 mM lidocaine ± cycloheximide, which blocks protein translation.^96^ Even with cycloheximide blockade, 10 mM lidocaine upregulated full-length caspase-3 and -7 proteins (Figures 6B and 6C).

**Figure 6.**
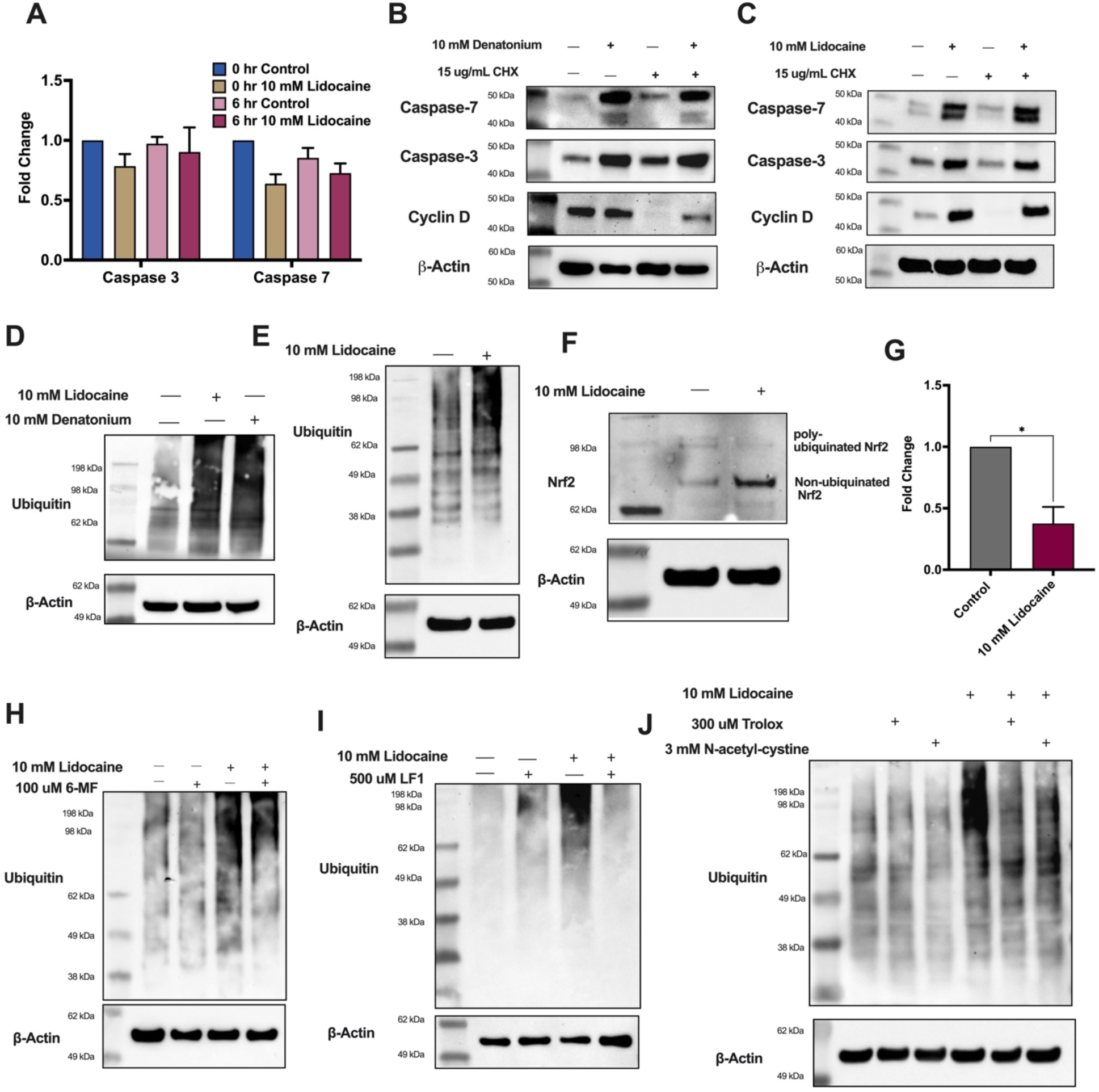
Lidocaine inhibits the proteasome via T2R14. **A)** SCC 47 caspase-3 (*CASP3)* and caspase-7 *(CASP7)* mRNA expression at 0 and 6 hours ± 10 mM lidocaine. Expression shown relative to 0-hour control. Mean +/- SEM with >3 separate cultures. No significance difference by paired t-test. **B-C)** SCC 47 cells were treated with (*B*)10 mM denatonium benzoate or (*C*) 10 mM lidocaine for 6 hours with or without 15 ug/mL cycloheximide (CHX). Caspase-3, caspase-7, cyclin D (cycloheximide protein synthesis control) and β-Actin primary antibody (1:1000) were used with goat anti-rabbit or anti-mouse IgG-horseradish peroxidase secondary antibodies (1:1000). **D-E)** SCC 47 cells (*D*) and FaDu cells (*E*) were treated with 10 mM lidocaine for 6 hours. Ubiquitin and β-Actin primary antibody (1:1000) were used with anti-rabbit or anti-mouse IgG-horseradish peroxidase secondary antibodies (1:1000). **F)** SCC 47 cells were treated with 10 mM lidocaine for 6 hours. Nrf2 and β-Actin primary antibody (1:1000) were used with anti-rabbit or anti-mouse IgG-horseradish peroxidase secondary antibodies (1:1000). **G)** mRNA expression of *FOXM1* in SCC 47 cells after treatment with 10 mM lidocaine for 6 hours. Expression is relative to control/untreated. Mean +/- SEM with >3 separate cultures. Significance determined by paired t-test. **(H-I)** SCC 47 cells were treated with 10 mM lidocaine ± 100 uM 6-MF (*H*) or 500 uM LF1 (*I*) for 6 hours. Ubiquitin and β-Actin primary antibody (1:1000) were used with anti-rabbit or anti-mouse IgG-horseradish peroxidase secondary antibodies (1:1000) Immunoblots shown are representative of 3 independent experiments using cells at different passage number on different days**. I)** SCC 47 cells were treated with 10 mM lidocaine ± 300 uM Trolox or 3 mM n-acetyl-L-cysteine for 6 hours. Ubiquitin and β-Actin primary antibody (1:1000) were used with anti-rabbit or anti-mouse IgG-horseradish peroxidase secondary antibodies (1:1000) All immunoblots shown are representative of ≥3 independent experiments using cells at different passage number on different days. P < 0.05 (*), P < 0.01 (**), P < 0.001 (***), and no statistical significance (ns or no indication).

The ubiquitin proteasome system tags unwanted or misfolded proteins for degradation.^97^ An accumulation of ubiquitin-tagged proteins indicates inhibition of the proteasome.^98^ We found that both lidocaine and denatonium cause an accumulation of poly-ubiquitinated proteins in SCC 47 and FaDu cells via Western blot (Figures 6D and 6E). To validate this observation, SCC 47 cells were incubated with 10 or 100 uM MG-132, a known proteasome inhibitor.^99^ MG-132 caused similar ubiquitin increases to those with 10 mM lidocaine or denatonium (Figure S8A). Lidocaine also downregulated the mRNA expression of *FOXM1,* a common a common hallmark of proteasome inhibition, and increased levels of Nrf-2, which is degraded by the proteasome at baseline (Figures 6F and 6G).^100,101^

Inhibition of T2R14 using 6-MF or LF1 reduced the accumulation of poly-ubiquinated proteins with lidocaine, linking T2R14 activation with proteasome inhibition (Figures 6H and 6I). When cells were treated with lidocaine in the presence of ROS inhibitors, Trolox or n-acetyl-n-cysteine, poly-ubiquinated protein accumulation was also rescued (Figure 6J). Together, this suggests that proteasome dysfunction may be a result of upstream mitochondrial dysfunction and excess ROS production caused by T2R14 activation.

To investigate the potential clinical applications of lidocaine, we used Matrigel to model a possible gel vehicle in which the lidocaine could be applied. Using both “thin” and “thick” formulations, we found that lidocaine induced cell death over 24 hours when compared to control via DIC images. The thick formulation appeared to have the most death, as compared to the thin formulation (Figure 7A). To test lidocaine in a 3D culture model, FaDu cells were grown in non-attached spheroids to model structure of HNSCC tumors. 10 mM lidocaine disbanded the spheroids as early as 8 hours and had a more profound effect at 18 hours (Figure 7B).

**Figure 7.**
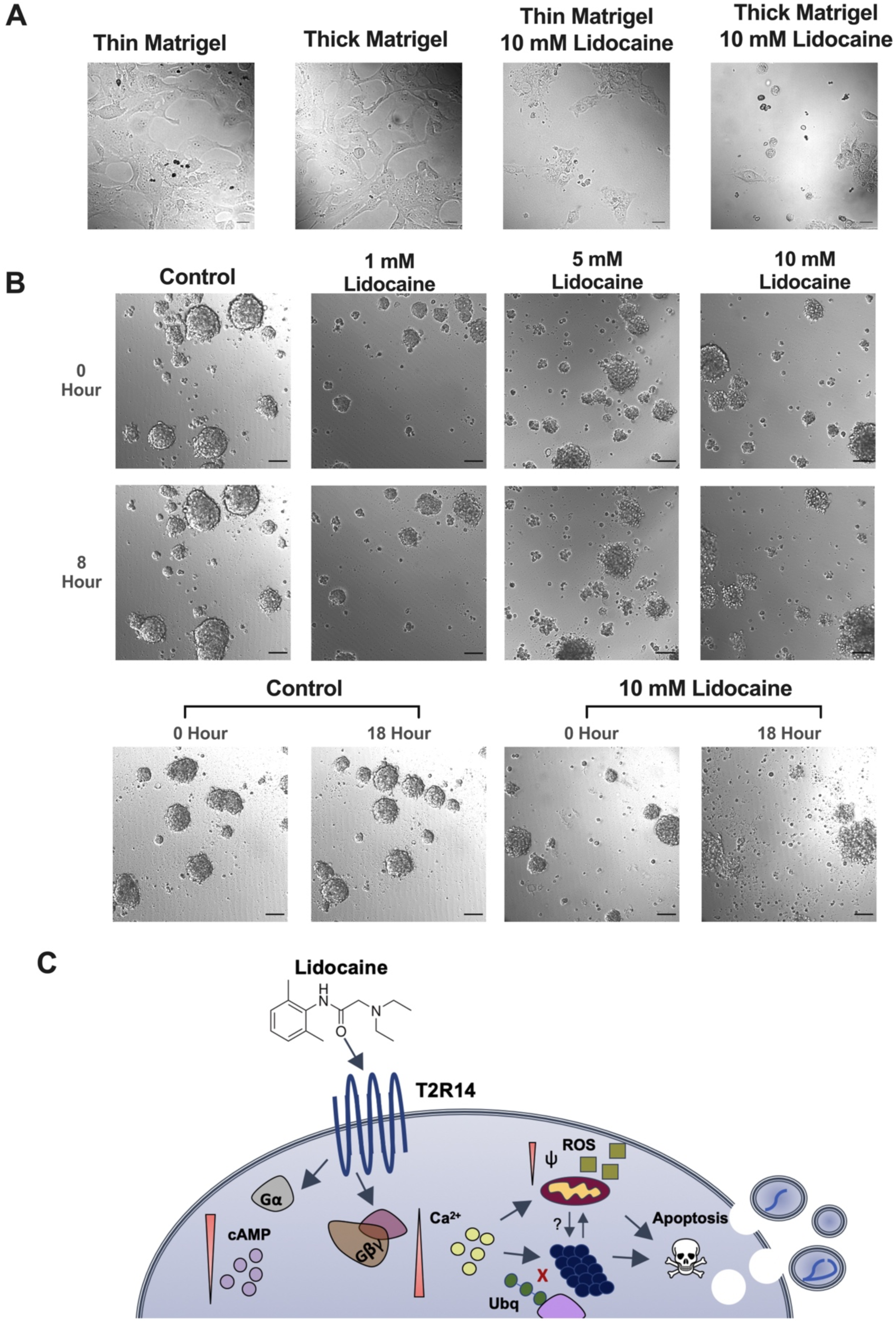
Matrigel model of lidocaine application. **A)** SCC 47 cells were subjected to Matrigel containing 10 mM lidocaine for 24 hours. DIC images of SCC 47 cells after 24 hours of lidocaine-Matrigel exposure with both thick and thin formulations. Scale bars = 30 µm. **B)** FaDu spheroids were grown for 5 days. They were incubated with 0 – 10 mM lidocaine for 8 or 18 hours. Scale bars = 30 µm. **C)** Working model of T2R14-induced apoptosis with lidocaine.

## Discussion

With roughly half of patients surviving past five years even with treatment, the outlook for HNSCC patients can be bleak.^7,102^ Standard treatment options (surgery, radiotherapy, chemotherapy) often leave lasting effects.^103–106^ Targeted therapies including immune checkpoint inhibitors (e.g. pembrolizumab and nivolumab)^1–4^ or EGFR inhibitors (e.g. cetuximab)^107,108^ have shown mixed results and are typically reserved for the recurrent and metastatic setting.^108^ Ultimately, the specificity of therapies for cancerous cells and not normal cells, remains as the most significant obstacle. Novel therapies or applications of existing therapies with favorable side-effect profiles must be discovered to improve outcomes for HNSCC patients.

Data here suggest that T2Rs may be attractive targets to leverage against HNSCCs using existing clinical compounds including the local anesthetic, T2R14-agonist, lidocaine. Although T2Rs are low affinity-GPCRs, their agonists have detrimental effects on HNSCC cellular function, mitochondrial membrane potential, and cell survival.^32^ In some cases of oral SCC, we found that T2R14 expression is upregulated, indicating that a personalized treatment approach based on receptor expression may be reasonable.^32^ T2R expression is not limited to just the oral cavity but also other epithelial regions affected by HNSCCs.^109^ An advantage of HNSCCs is that many of the affected mucosal sites are easily accessible, and thus bitter agonists like lidocaine could be implemented into clinical practice as topical anticancer therapies (e.g., an eluting gel or topical rinse).^110^ With high solubility, documented safety data, and beneficial palliative effects, lidocaine has the potential to be an easily translatable treatment.^111,112^

Here, we established interactions between lidocaine and T2R14 by recording Ca^2+^ responses in HNSCC immortalized cells lines. Compared to other T2R agonists including flufenamic acid, thujone, and denatonium benzoate (one of the most bitter compounds known in terms of taste perception on the tongue), lidocaine stimulation produced the largest Ca^2+^ responses, likely because it potently and specifically activates T2R14. Procaine which has a bitter taste but is not known to activate T2R14, had minimal effects on Ca^2+^, suggesting the effects of lidocaine are not due to Na^+^ channel inhibition. In addition, ATP produced significant Ca^2+^ responses in HNSCC cells at only 100 uM yet did not reduce cell viability like bitter agonists (Figures 1E-1G; Figure S6D).^51^

Both suramin and pertussis toxin dampened lidocaine Ca^2+^ responses, indicative of GPCR activation (Figures 3A and B).^66,67^ However, suramin and pertussis toxin did not fully abolish the Ca^2+^ response with lidocaine. GPCRs are known to localize to the cellular membrane, however, immunofluorescent staining of T2R14 revealed cytoplasmic localization (Figure 3L). Some GPCRs do localize in cytoplasm or to the nuclear membrane, and this difference may affect the affinity of both G-protein inhibitors.^113^ Pertussis toxin did not affect Ca^2+^ response with denatonium benzoate, suggesting that the compound interacts with different T2Rs that are differently coupled (Figure S2B).^114,115^ Ca^2+^ responses with denatonium benzoate, but not lidocaine, were dampened with YM-254890, a Gα_q_ inhibitor, and with shRNA knockdown of T2R4 (Figure S2C-S2E).^116^ Although T2Rs are primarily coupled to Gα_gustducin_ or Gα_i_, there is also evidence that they may be coupled to Gα*_q_*_14_ as well.^117^ This could explain the dampened Ca^2+^ response with YM-254890 with denatonium, but not with lidocaine. This further supports data that shows lidocaine interacts with different T2Rs than denatonium.

Lidocaine decreased cAMP and increased DAG, secondary messengers of GPCR activation (Figures 3C and 3D). We found that 6-MF, LF1, and LF22, specific T2R14 antagonists, also dampened Ca^2+^ responses with lidocaine and other T2R14 agonists, suggesting activation of T2R14 (Figures 3E-3J).^35^ Similar to suramin and pertussis toxin, the Ca^2+^ responses with 6-MF were not completely blocked, possibly due to incomplete antagonism or cross-reactivity with another T2R. However, LF1 and LF22 dampened Ca^2+^ response with lidocaine more robustly. LF1 and LF22 are derivatives of flufenamic acid, as they share the same binding pocket within T2R14. This suggests that lidocaine may share a binding site with flufenamic acid and that LF1 and LF22 disrupt these binding sites better than 6-MF.

Studies have investigated whether lidocaine can be repurposed as a therapeutic in several cancers.^38,42,118,119^ This is a promising and low-cost endeavor as the local anesthetic is already FDA approved.^120^ However, none of these studies have focused on HNSCCs or have considered activation of T2R14.^86,121^ Here, we found that lidocaine alone significantly decreased HNSCC cell viability and depolarized the mitochondrial membrane (Figures 4A-4D). Lidocaine decreased cell proliferation at just 6 hours. This corroborates the observed decrease in mitochondrial membrane potential. The mitochondria are a primary source of NADH production, which is the parameter in which cell viability is measured with XTT. Complimentary to this, lidocaine also caused the production of ROS, further indicating mitochondrial dysfunction and cellular stress (Figures 4F and 4G). Lidocaine induced apoptosis in HNSCC cells and 6-MF blocked this observed response, linking T2R14 activation with apoptotic induction (Figure 5).^90,122,123^

Our data also suggest lidocaine inhibits proteasomal degradation of several proteins and causes accumulation of poly-ubiquinated proteins (Figure 6).^124,125^ The proteasome complex made up of subunits that have protease activity. This complex can capture poly-ubiquinated proteins, unfold them, and subject them to proteolysis.^126^ When this system is inhibited, proteins that are meant for degradation accumulate. The accumulation can induce apoptosis. Lidocaine also downregulated *FOXM1* and activated Nrf2, which are indicative of proteasome inhibition (Figures 6F and 6G). *FOXM1* and Nrf2 can also be affected in the same ways when there is oxidative stress, as observed here with mitochondrial ROS production (Figure 4E and 4F).^127,128^ The UPS can be negatively regulated by the mitochondrial via increased ROS and decreased ATP.^129^ Lidocaine increases ROS and we observed that proteasome function rescue when ROS are inhibited in the presence of lidocaine (Figure 6J). In addition, when T2R14 is antagonized, proteasome inhibition caused by lidocaine is rescued (Figures 6I and 6J). This suggests that proteasome inhibition is a result of T2R14 activation with lidocaine. This regulation may be a downstream effect of mitochondrial dysfunction and ROS caused by lidocaine as it is rescued by two structurally distinct antioxidants.

MG-132, a known proteasome inhibitor, caused poly-ubiquinated protein accumulation similar to lidocaine, however, MG-132 does not induce apoptosis within the timeframe of the induction with lidocaine (Figures S8A and S8B). This indicates that there may be other underlying pro-apoptotic mechanisms that could be independent or influence/influenced by proteasome inhibition, such as the discussed mitochondrial dysfunction. Whether or not this is a lidocaine-T2R14 specific response or if this is a global response with all T2R14 agonists is also unknown.

It is also possible that, independent of T2R14 activation, lidocaine is able to diffuse into the cell.^130^ Lidocaine has been suggested to inhibit the proteasome by binding to the β5 subunit of the 20s proteasome in p53-postive cancer cells in *in vitro* assays.^131^ This may contribute to apoptosis observed with lidocaine stimulation, either downstream of T2R14 activation or independently. However, our data suggest that the mechanism of lidocaine proteasome inhibition in intact cells instead depends on T2R14 signaling.

It is highly important to continue to study and uncover new therapeutics and therapeutic targets to improve HSNCC patient outcomes. Our data suggest using lidocaine to treat HNSCCs (beyond its existing use in pain relief) could prove to be a promising strategy, warranting further *in vivo* and clinical investigation. Drugs that inhibit the catalytic activity of the proteasome, including hallmark drugs like bortezomib and ixazomib are used as chemotherapeutics in select cancers, taking advantage of the high rates of protein-turnover in malignancy.^132^ It’s important to note that lidocaine kills both FaDu (-HPV) and SCC47 (+HPV) cells. Lidocaine may be a potent therapy for both HPV positive and negative HNSCCs. Not all treatments are effective for both types of HNSCCs. Due to accessibility of head and neck anatomic sites, lidocaine could potentially be delivered as a topical cream or local injection independently or in conjunction with existing treatment approaches. Specifically, lidocaine may be most useful in the subset of patients with elevation of T2R14 in tumor tissue as a component of a “personalized medicine” cancer treatment regimen.^32^

## Acknowledgements

We thank M. Victoria (University of Pennsylvania) for excellent technical assistance. We thank N. Cohen (University of Pennsylvania) for helpful discussions. This study was supported by T32GM008076 (Z.A.M.), an American Head and Neck Society Pilot Grant (R.M.C.), R01DC016309 (R.J.L.), R01AI167971 (R.J.L.), and R21DC020041 (R.J.L.).

## CRediT authorship contribution statement

Zoey A. Miller: Conceptualization, Investigation, Methodology, Formal analysis, Writing – original draft, Writing – review & editing; Jennifer F. Jolivert: Investigation; Arielle Mueller: Investigation; Ray Ma: Investigation; Sahil Muthuswami: Investigation; April Park: Investigation; Derek McMahon: Conceptualization, Methodology; Ryan M. Carey: Resources, Conceptualization, Funding acquisition; Writing-review & editing; Robert J. Lee: Conceptualization, Writing – review & editing, Project administration, Supervision, Funding acquisition.

## Declaration of interests

The authors declare no competing interests.

## STAR ★ METHODS

### KEY RESOURCES TABLE

| REAGENT OR RESOURCE | SOURCE | IDENTIFIER |
| --- | --- | --- |
| **Antibodies** |  |  |
| Anti-caspase-3 (D3R6Y) | Cell Signaling | 9662 |
| Anti-caspase-7 (D2Q3L) | Cell Signaling | 9492 |
| Anti-cyclin D | Cell Signaling | 2978 |
| Anti-TAS2R14/TRB1 | LS Bio | C386266 |
| Anti-ubiquitin (P4D1) | Cell Signaling | 3936 |
| Anti-nrf2 | Abcam | Ab62352 |
| Anti-Bax | Cell Signaling | 2772 |
| Anti-β-actin | Cell Signaling | 3700 |
| Anti-rabbit AlexaFluor 647 | Life Technologies | A31537 |
| Anti-rabbit IgG, HRP Conjugated | Santa Cruz | SC-2004 |
| Anti-mouse IgG, HRP Conjugated | Santa Cruz | SC-2005 |
| **Bacterial and Virus Strains** |  |  |
| Red Upward DAG sensor Bacman | Montana Molecular | U0300R |
| **Chemicals** |  |  |
| Lidocaine | Cayman Chemical | 20081 |
| Nigericin (sodium salt) | Cayman Chemical | 11437 |
| Ryanodine | Cayman Chemical | 14962 |
| Suramin | Cayman Chemical | 11126 |
| Isoproterenol | Cayman Chemical | 15592 |
| YM254890 | Cayman Chemical | 29735 |
| Trolox | Cayman Chemicals | 10011659 |
| N-acetyl-L-Cysteine | Cayman Chemicals | 20261 |
| Pertussis Toxin | Tocris | 3097 |
| Denatonium Benzoate | TCI Chemicals | D2124 |
| ATP | Millipore Sigma | A9187 |
| (-)-α-Thujone | Millipore Sigma | 89231 |
| Flufenamic acid (FFA) | Millipore Sigma | F9005 |
| Saponin | Millipore Sigma | S7900 |
| Bovine Serum Albumin | Millipore Sigma | A2153 |
| Caffeine | Millipore Sigma | 58-08-2 |
| Crystal Violet | Millipore Sigma | C0775-25G |
| JC-1 | ThermoFisher | T3168 |
| CellEvent™ Caspase-3/7 Detection Reagent | ThermoFisher | C10423 |
| XTT (sodium 3'-(1- (phenylaminocarbonyl)- 3,4-tetrazolium)-bis (4-methoxy6-nitro) benzene sulfonic acid hydrate | ThermoFisher | X6493 |
| Lipofectamine 3000 | ThermoFisher | L3000075 |
| Fluo-4, AM, cell permeant | ThermoFisher | F14201 |
| 6-methoxyflavanone | ThermoFisher | 11420768 |
| TriZol | ThermoFisher | 15596026 |
| Normal Donkey Serum | Abcam | ab7475 |
| Fluoroshield Mounting Medium w/ DAPI | Abcam | Ab104131 |
| LF1 | Enamine | Z85879385 |
| LF22 | Enamine | Z3906872218 |
| Phalloidin AlexaFluor 488 | Life Technologies | A12379 |
| Matrigel Matrix | Corning | 356255 |
| **Critical Commercial Assays** |  |  |
| High-Capacity cDNA Reverse Transcription Kit | ThermoFisher | 4368814 |
| DC Protein Assay Kit | BioRad | 5000111 |
| Direct-zol RNA Miniprep | Zymo Research | R2051 |
| SuperSignal West Pico PLUS | ThermoFisher | 34577 |
| Clarity Max Western ECL Substrate | BioRad | 1705060 |
| **Experimental Models** |  |  |
| SCC 4 | ATCC | CRL-1624 |
| SCC 90 | ATCC | CRL-3239 |
| UMSCC 47 (SCC 47) | Millipore Sigma | SCC071 |
| HEK293T | ATCC | CRL-11268 |
| RPMI 2650 | ATCC | CCL-30 |
| NCI H520 | ATCC | HTB-182 |
| **Oligonucleotides** |  |  |
| Taqman Primers for RyR1 | ThermoFisher | Hs0016699_m1 |
| Taqman Primers for RyR2 | ThermoFisher | Hs00181461_m1 |
| Taqman Primers for RyR3 | ThermoFisher | Hs00168821_m1 |
| Taqman Primers for IP3R1 | ThermoFisher | Hs00181881_m1 |
| Taqman Primers for IP3R2 | ThermoFisher | Hs00181916_m1 |
| Taqman Primers for IP3R3 | ThermoFisher | Hs01573539_m1 |
| Taqman Primers for TAS2R4 | ThermoFisher | Hs00249646_s1 |
| Taqman Primers for TAS2R14 | ThermoFisher | Hs00256800_s1 |
| Taqman Primers for BAX | ThermoFisher | Hs00180269_m1 |
| Taqman Primers for BAD | ThermoFisher | Hs00188930_m1 |
| Taqman Primers for Bcl-2 | ThermoFisher | Hs04986394_s1 |
| Taqman Primers for Caspase-3 | ThermoFisher | Hs00234387_m1 |
| Taqman Primers for Caspase-7 | ThermoFisher | Hs00169152_m1 |
| Taqman Primers for FOXM1 | ThermoFisher | Hs01073586_m1 |
| Taqman Primers for UBC | ThermoFisher | Hs01871556_s1 |
| **Recombinant DNA** |  |  |
| 4mtD3cpv | Addgene | 36324 |
| Flamindo2 | Addgene | 73938 |
| D1ER | Addgene | 36325 |
| pRS hTAS2R4 shRNA | Origene | TR30123B |
| pRS hTAS2R14 shRNA | Origene | TR301238A |
| **Software and Algorithms** |  |  |
| MetaFluor | Molecular Devices | N/A |
| MetaMorph | Molecular Devices | N/A |
| QuantStudio 5 | Applied Biosystems, Inc | N/A |
| Prism v9 | GraphPad Software | N/A |
| ImageJ/FIJI | Open Source<sup>133</sup> | N/A |

### RESOURCE AVAILABILITY

#### Lead contact

Robert J. Lee.

#### Materials availability

Further information and requests for resources and reagents should be directed to and will be fulfilled by Robert J. Lee. Vectors or cell lines generated for these studies will be provided upon request.

#### Data and Code availability

The published article includes all datasets generated or analyzed during this study. Raw numerical values used to generate bar graphs or traces are available upon request.

### METHODS

#### Cell Culture

SCC4 (ATCC CRL-1624), SCC47 (UM-SCC-47; Millipore SCC071), SCC90 (ATCC CRL-3239), FaDu (ATCC HTB-43), RPMI 2650 (ATCC CCL-30), and HEK 293T (ATCC CRL-3216) cell lines were from ATCC (Manassas, VA, USA) or MilliporeSigma (St. Louis, MO USA). All cell lines were grown in submersion in high glucose Dulbecco’s modified Eagle’s medium (Corning; Glendale, AZ, USA) with 10% FBS (Genesee Scientific; El Cajon, CA, USA), penicillin/streptomycin mix (Gibco; Gaithersburg, MD, USA), and nonessential amino acids (Gibco). Stable SCC90 T2R4 shRNA knockdown cells were generated previously.^32^ Transient HEK-293T T2R14 shRNA knockdown cells were produced using lipofectamine 3000 reagents.

#### Quantitative reverse transcription PCR (qPCR)

Cell cultures were resuspended in TRIzol (Thermo Fisher Scientific; Waltham, MA, USA). RNA was isolated and purified (Direct-zol RNA kit; Zymo Research), reverse transcribed via High-Capacity cDNA Reverse Transcription Kit (Thermo Fisher Scientific), and quantified using TaqMan qPCR probes for *TAS2R4, TAS2R14, CASP3, CASP7, RyR1, RyR2, RyR3, IP3R1, IP3R2, IP3R3, FOXM1,* and *UBC* (QuantStudio 5; Thermo Fisher Scientific). *UBC* was used as an endogenous control due to its stability in expression in cancer cells.^134^

#### Live Cell Imaging

For calcium (Ca^2+^) imaging, cells were loaded with 5 µM of Fluo-4-AM (Thermo Fisher Scientific) for 50 min at room temperature in the dark. Hank’s Balanced Salt Solution (HBSS) buffered with 20 mM HEPES (pH 7.4) was used as an imaging buffer, containing 1.8 mM Ca^2+^. Cells were imaged using a Nikon Eclipse TS100 (20x 0.75 NA PlanApo objective), FITC filters (Chroma Technologies), QImaging Retiga R1 camera (Teledyne; Tucson, AZ, USA), MicroManager, and Xcite 120 LED Boost (Excelitas Technologies; Mississauga, Canada). For experiments in Ca^2+^-free (0-Ca^2+^) HBSS, cells were loaded with Fluo-4-AM as described above in normal Ca^2+^-containing HBSS. Bitter agonists were dissolved in HBSS with no added Ca^2+^ plus 10 mM EGTA (0-Ca^2+^ HBSS) and was used in the place of regular HBSS (no EGTA). At the start of the experiment, 300 uL of bitter agonist in 0-Ca^2+^ HBSS was added to 60 uL of Ca^2+^-containing HBSS for final concentrations of ∼8.3 mM EGTA and 0.3 mM Ca^2+^ with a predicted free extracellular Ca^2+^ of 2.1×10^-9^ mM (MaxChelator, Chris Patton, Stanford University, https://somapp.ucdmc.ucdavis.edu/pharmacology/bers/maxchelator/). This method resulted in an acute removal of extracellular free Ca^2+^ at the exact time of agonist addition.

For mitochondrial and endoplasmic reticulum Ca^2+^ imaging, pcDNA-4mt3cpv or pcDNA-D1ER was transfected into cells using lipofectamine 3000 reagents 24-48 hours prior to imaging.^60^ Live cell images were taken on Olympus IX-83 microscope (20x 0.75 NA PlanApo objective), CFP/YFP filters (Chroma 89002-ET-ECFP/ EYFP) in excitation and emission filter wheels (Sutter Lambda LS), Orca Flash 4.0 sCMOS camera (Hamamatsu, Tokyo, Japan), Meta-Fluor (Molecular Devices, Sunnyvale, CA USA), and XCite 120 LED Boost (Excelitas Technologies). For cyclic adenosine monophosphate (cAMP) imaging, pcDNA3.1(-)Flamindo2 was transfected into cells as described above 48 hours prior to imaging.^70^ Cells were imaged as described above using FITC filters.

For diacylglycerol (DAG) imaging, RED Upward DAG Assay Kit (Montana Molecular; Bozeman, Montana, USA) was transduced into cells using 2x the recommended reagents. Cells were used 24 hours-post transduction and imaged as described above using TRITC filters.

Novel T2R14 inhibitors LF1 and LF22 were identified based on a structure-search using information from and purchased from a chemical vendor (Enamine, Monmouth Jct., NJ; cat # Z85879385 and Z3906872216). Compounds were dissolved in DMSO at 500 mM and used at 1:1000 as indicated. Stock solutions were stored at -20° C.^135^

For superoxide imaging, MitoSox fluorescent dye (Thermo Fisher Scientific) was used as described by the supplier following bitter agonist exposure.

#### Cell Viability and Proliferation

Cells were incubated with bitter agonists for 6 or 24 hours at 37 °C. Crystal violet (0.1% in deionized water with 10% acetic acid) was used to stain remaining adherent cells. Stains were washed with deionized water and left to dry at room temperature. Stains were dissolved with 30% acetic acid in deionized water. Absorbance values at 590 nm were measured.

XTT dye and phenazine methosulfate were added to cells in combination with bitter agonists. Measurements were taken at 475 and 660 nm over 6 hours. Absorbance values were measured on a Tecan (Spark 10M; Mannedorf, Switzerland).

Lidocaine was dissolved in Matrigel (Corning) on ice. Thin formulation was made by combining 75 µL Matrigel and 75 µL of 20 mM lidocaine in DMEM. Thick formulation was made by combining 150 µL Matrigel and 150 µL of 20 mM lidocaine dissolved in Dulbecco’s modified Eagle’s medium (Corning; Glendale, AZ, USA) with 10% FBS (Genesee Scientific; El Cajon, CA, USA), penicillin/streptomycin mix (Gibco; Gaithersburg, MD, USA), and nonessential amino acids (Gibco). The lidocaine Matrigel was added on top of cells in 8-well glass chamber slides (∼0.86 cm^2^ surface area per well). After 24 hours, live cell images were taken on Olympus IX-83 microscope (20x 0.75 NA PlanApo objective), Orca Flash 4.0 sCMOS camera (Hamamatsu, Tokyo, Japan), Meta-Fluor (Molecular Devices, Sunnyvale, CA USA), and Differential Interference Contrast (DIC) filter.

FaDu cells were grown into tumor spheroids on low-attachment 96-well plates using F-12 media supplemented as previously described.^136^ Spheroids were cultured for 5 days prior to exposure to lidocaine. Cells were imaged using the same technique described above for Matrigel.

#### Mitochondrial membrane potential and apoptosis measurements

Cells were loaded with JC-1 dye for 15 minutes. Bitter agonists were added as indicated. Fluorescence was measured at 488nm excitation and 535 (green) and 590 (red) nm emission. Live cell images of the assay were taken on Olympus IX-83 microscope (20x 0.75 NA PlanApo objective), FITC and TRITC filters (Chroma Technologies), Orca Flash 4.0 sCMOS camera (Hamamatsu, Tokyo, Japan), Meta-Fluor (Molecular Devices, Sunnyvale, CA USA), and XCite 120 LED Boost (Excelitas Technologies). Quantitative fluorescence values were measured on a Tecan Spark 10M.

CellEvent Caspase 3/7 dye was added to cells in combination with bitter agonists. Fluorescence was measured at 495 nm excitation and 540 nm emission. Live cell images were taken as described above for JC-1 assays (only FITC filter). Quantitative fluorescence values were measured on a Tecan Spark 10M.

#### Western Blotting

Protein was harvested using lysis buffer (10 mM Tris pH 7.5, 150 mM NaCl, 10 mM KCl, 0.5% deoxycholate, 0.5% Tween, 0.5% IGePawl, 0.1% SDS). Protein content was quantified using DC Protein Assay (BioRad; Hercules, CA, USA). 50 – 80 µg of protein were loaded into gel (Bis-Tris 4-12%, 1.5 mm) with 4x loading buffer (200 mM Tris pH 6.8, 40% glycerol, 8% SDS, 0.1% Bromophenol Blue), and 5% [3-mercaptoethanol. Gel was run using MES running buffer (Tris Base 50 mM pH 7.3, MES 50 mM, SDS 0.1%, EDTA 1 mM). Novex Sharp Pre-Stained Protein Standard, SeeBlue Plus2 Pre-stained Protein Standard (Thermo Fisher), or Odyssey One-Color Protein Molecular Weight Marker (LI-COR) were used as molecular markers. Gel was transferred using Bis-Tris transfer buffer (25 mM bicine, 25 mM bis-tris, 1 mM EDTA, 10% methanol). Blots were blocked for 1 hour in TBST (24.8 mM tris acid & base, 1.5 NaCl, 0.5% Tween-20) with 5% milk. Primary antibodies were used 1:1000 or 1:500 with HRP-conjugated chemiluminescent secondary antibodies 1:1000 in TBST with 5% BSA (primary) or milk (secondary). Blots were imaged using Clarity Max Western ECL Substrate (BioRad) using ChemDoc MP Imaging System (BioRad).

#### Immunofluorescence

Cells were fixed on glass with 4% formaldehyde in PBS w/ Ca^2+^ and Mg^2+^. Cells were blocked and permeabilized in blocking buffer containing PBS with Ca^2+^ and Mg^2+^, 0.2% saponin, 3% normal donkey serum, 1% BSA and 0.1% Triton X-100. Cells were incubated in primary antibody at either 1:100 or 1:200 overnight. Secondary antibody incubation was for 2 hours at 1:500. Phalloidin was used to stain actin at 1:400 and DAPI was used to stain the nucleus. Images were taken on Olympus Live Cell Imaging System (system as described in live cell imaging) with 60x oil objective.

#### Statistical Analysis

Data were analyzed using *t-*test (two comparisons only) or one-way ANOVA (>2 comparisons) in GraphPad Prism (San Diego, CA, USA). Both paired and unpaired *t*-test were used when appropriate. Bonferroni and Dunnett’s posttest for one-way ANOVA were used when necessary. All figures used the following annotations: P < 0.05 (*), P < 0.01 (**), P < 0.001 (***), and no statistical significance (ns). All data points represent the mean +/- SEM.

## Notes

### Competing Interest Statement

The authors have declared no competing interest.

### Summary of Updates

This version of the manuscript has been revised to include new data and text in both the main paper and supplement.

