## Supplemental Figure for "Lidocaine Induces Apoptosis in Head and Neck Squamous Cell Carcinoma Through Activation of Bitter Taste Receptor T2R14"

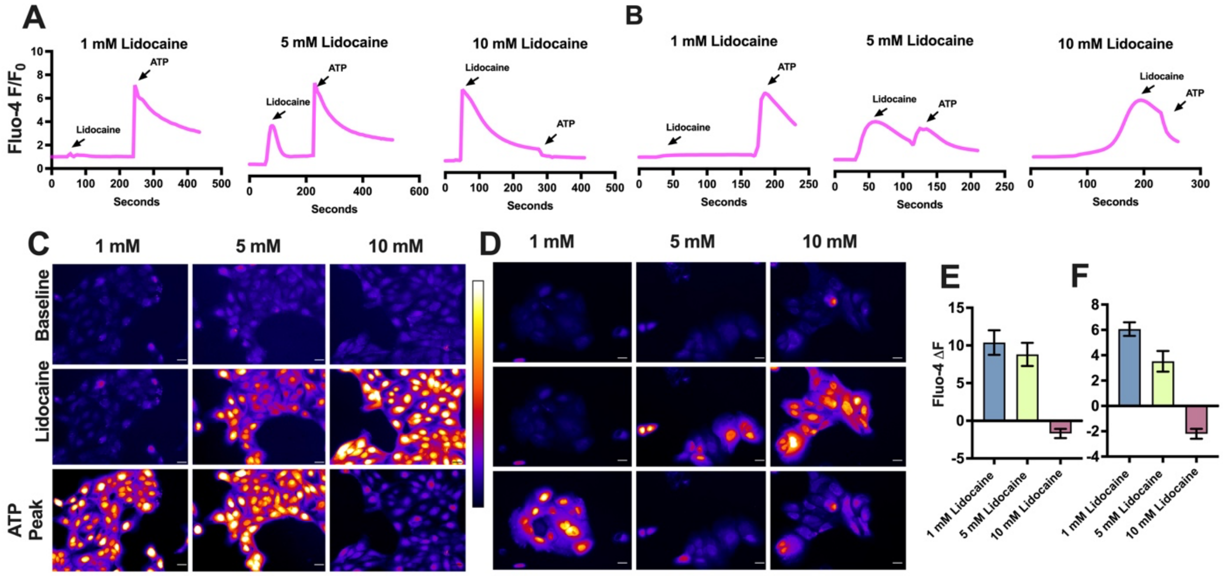


**Supplemental Figure 1. Lidocaine exhausts purinergic $Ca^{2+}$ responses in HNSCC cells.** SCC 47 and SCC 4 cells were loaded with Fluo-4 and imaged for subsequent $Ca^{2+}$ responses with 1 – 10 mM lidocaine followed by 100 μM ATP. **A)** SCC 47 fluorescent $Ca^{2+}$ responses over time and representative images of responses of baseline, lidocaine peak and ATP peak with 1 – 10 mM lidocaine followed by 100 μM ATP. Scale bars = 30 μm. **B)** SCC 4 fluorescent $Ca^{2+}$ responses over time and representative images of responses of baseline, lidocaine peak and ATP peak with 1 – 10 mM lidocaine followed by 100 μM ATP. Scale bars = 30 μm. **C-D)** Representative images of peak $Ca^{2+}$ responses described in traces above for **C)** SCC 47 and **D)** SCC 4. **E)** SCC 47 and **F)** SCC 4 changes in fluorescence with secondary ATP simulation.

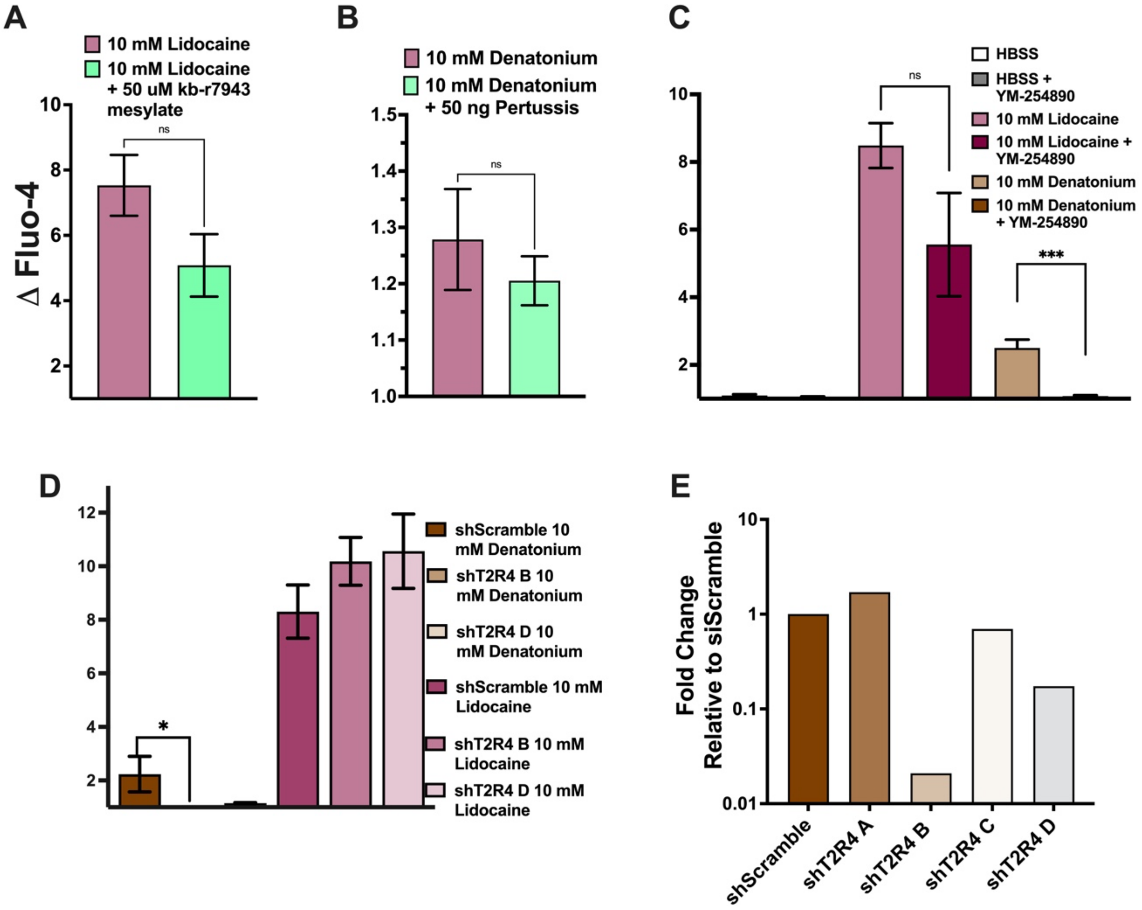


**Supplemental Figure 2. Lidocaine activates separate intracellular $Ca^{2+}$ pathway than denatonium benzoate.** HNSSC cell lines, SCC 47 and SCC 90, were loaded with Fluo-4 and imaged for subsequent $Ca^{2+}$ responses with 10 mM lidocaine or 10 mM denatonium. **A)** SCC 47 peak fluorescent $Ca^{2+}$ response with 10 mM lidocaine with or without prior 2-minute stimulation with 50 μM kb-r7943. **B)** SCC 47 peak fluorescent $Ca^{2+}$ response with 10 mM denatonium with or without prior 18-hour pertussis toxin incubation. Peak fluorescent mean +/- SEM with >3 separate cultures. Significance by unpaired t-test. **C)** SCC 47 peak fluorescent $Ca^{2+}$ responses with HBSS, 10 mM lidocaine, or 10 mM denatonium benzoate with or without prior 1-hour incubation with 1 μM YM-254890. Peak fluorescent mean +/- SEM with >3 separate cultures. Significance by unpaired t-test between bitter agonist response and bitter agonist response with YM-254890. **D)** SCC 90 siT2R4 peak fluorescent $Ca^{2+}$ responses with 10 mM lidocaine or 10 mM denatonium. Peak fluorescence mean +/- SEM with >3 experiments using separate cultures. Significance by 1-way ANOVA with Bonferroni posttest comparing shScramble response with 10 mM lidocaine or denatonium to each shT2R4 B or D with respective bitter agonist. **E)** *Tas2R4* mRNA expression in SCC 90 siT2R4 cells relative to shScramble.

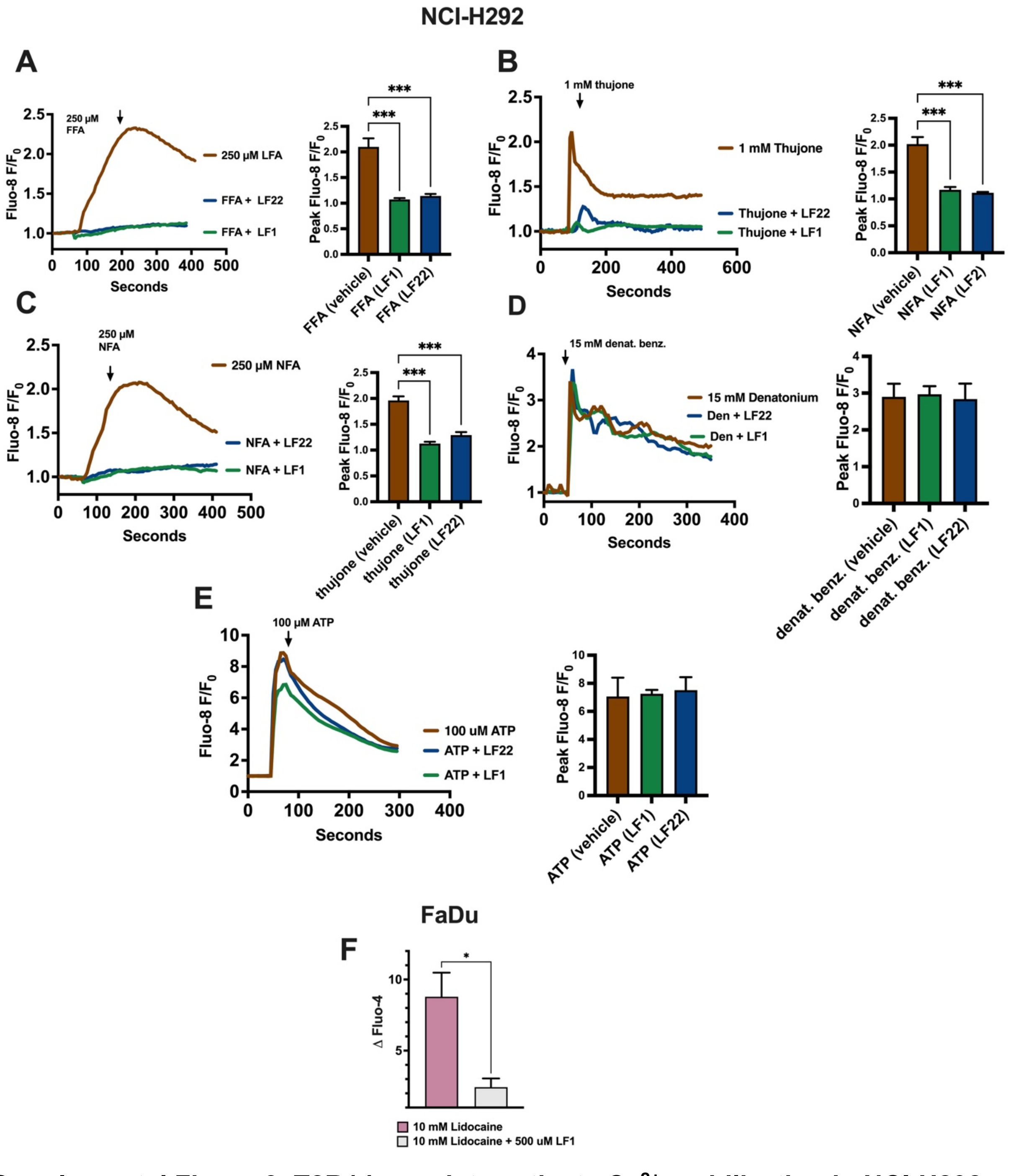


**Supplemental Figure 3. T2R14 agonists activate $Ca^{2+}$ mobilization in NCI-H292 cells. A-E)** NCI-H292 cells were loaded with Fluo-4 and imaged for subsequent $Ca^{2+}$ responses with *A)* flufenamic acid (FFA), *B)* thujone, *C)* niflumic acid (NFA), *D)* denatonium benzoate, or *E)* ATP. Bar graphs represent peak $Ca^{2+}$ responses +/- prior 1-hour incubation with LF1 or LF22. Peak fluorescence mean +/- SEM with >3 experiments using separate cultures. Significance by 1-way ANOVA with Bonferroni posttest comparing agonist alone with agonist + LF1 or LF22. **F)** FaDu peak fluorescent $Ca^{2+}$ response with lidocaine +/- 1-hour prior incubation with 500 uM LF1. Significance determined by unpaired t-test. $P < 0.05$ (*), $P < 0.01$ (**), $P < 0.001$ (***), and no statistical significance (ns or no indication).

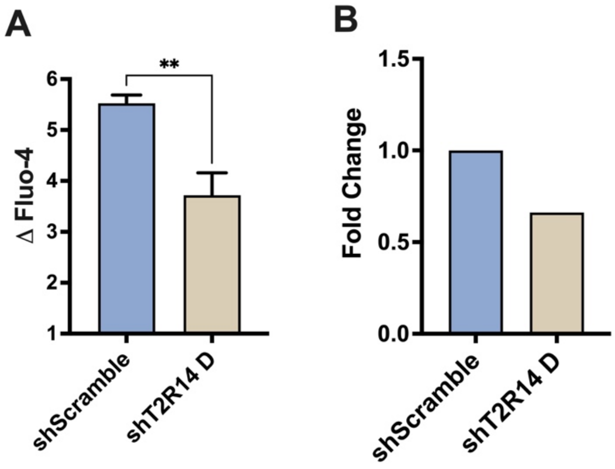


**Supplemental Figure 4. Silencing T2R14 with shRNA dampens $Ca^{2+}$ response in HEK-293T cells. A)** HEK-293T shScramble or shT2R14 peak $Ca^{2+}$ responses with lidocaine. Significance determined by unpaired t-test. **B)** HEK-293T mRNA *Tas2R14* expression normalized to shScramble. Significance determined by unpaired t-test. $P < 0.05$ (*), $P < 0.01$ (**), $P < 0.001$ (***), and no statistical significance (ns or no indication).

A

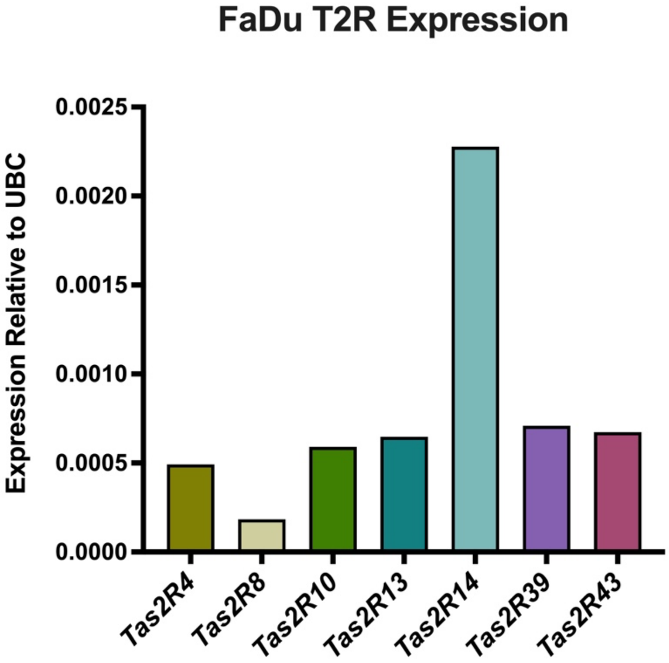


B

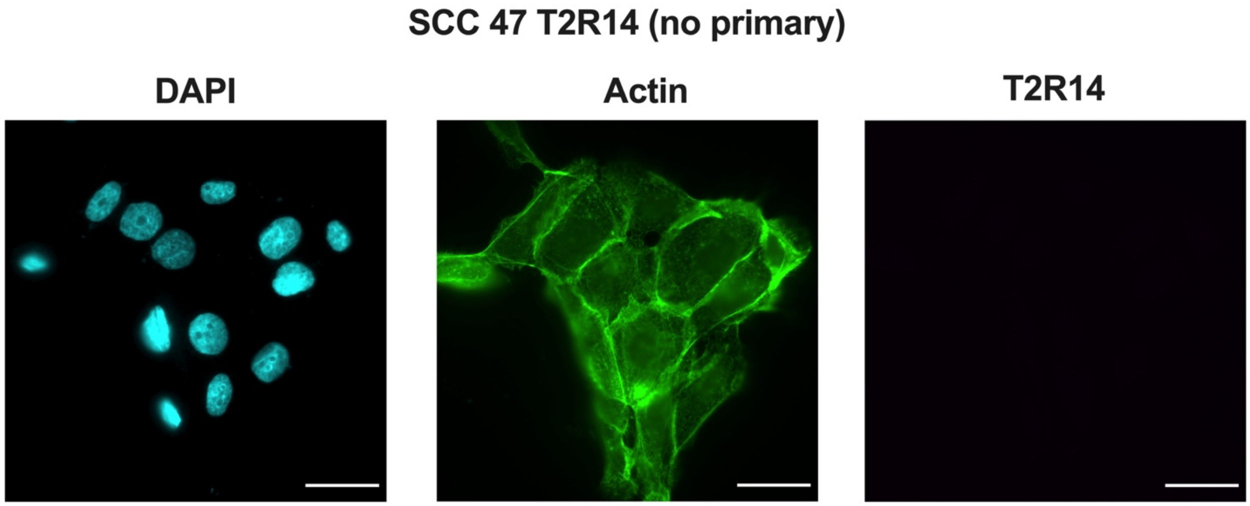


**Supplemental Figure 5. FaDu cells express T2R14. A)** mRNA expression of T2R4, T2R8, T2R10, T2R13, T2R14, T2R39, and T2R43 in FaDu cells. Expression relative to UBC, endogenous control. **B)** T2R14 control expression in SCC 47 via immunofluorescence stain with DAPI (nucleus) and phalloidin (actin) (no primary antibody).

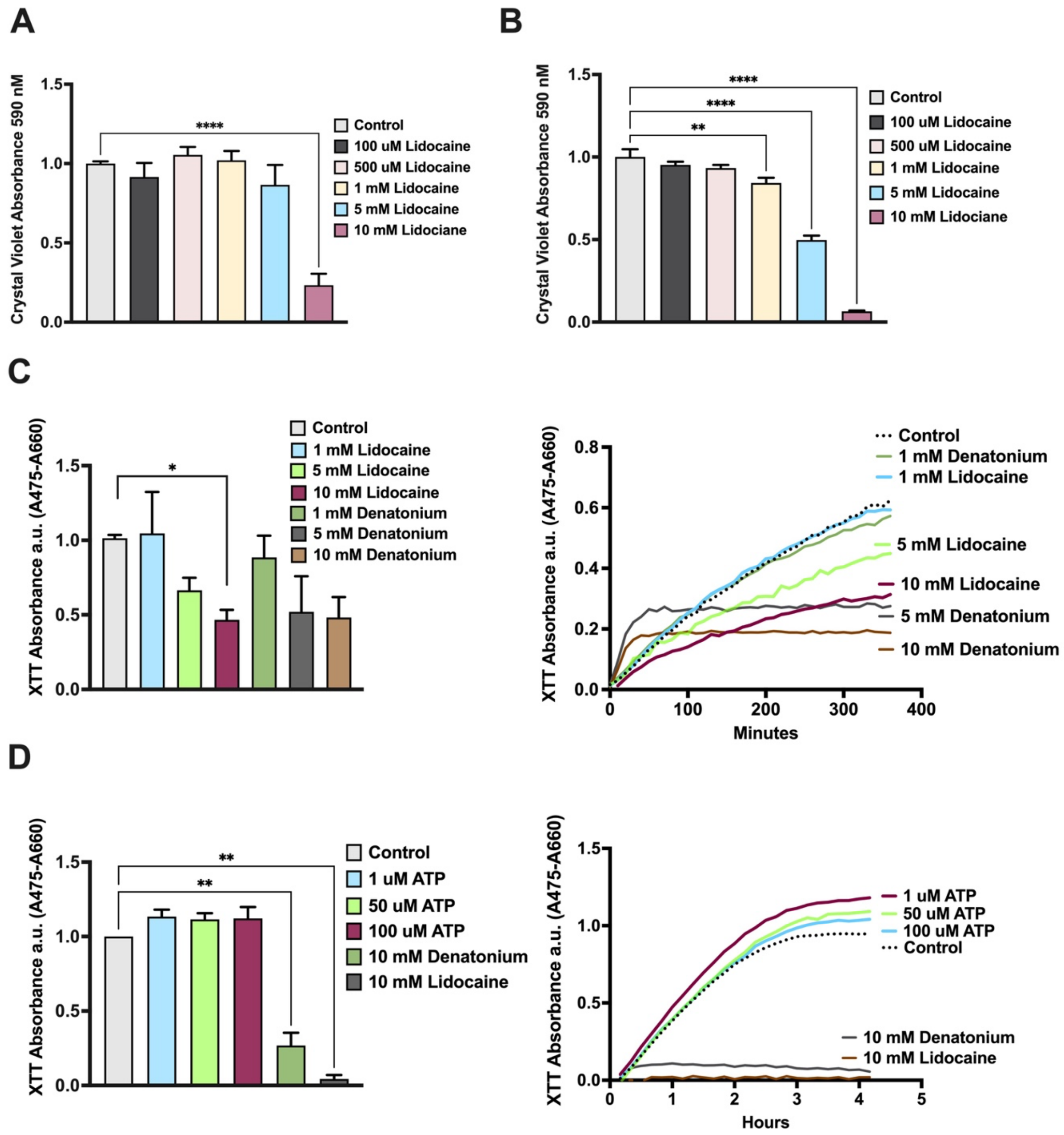


**Supplemental Figure 6. Lidocaine decreases cell viability and proliferation. A)** SCC 47 cells were incubated with 0 – 10 mM lidocaine for 6 or 24 hours and stained with crystal violet to determine cell proliferation. SCC 47 crystal violet stain absorbance values (590 nm) after 6 hours with 0 – 10 mM lidocaine. **B)** SCC 47 crystal violet stain absorbance values (590 nm) after 24 hours with 0 - 10 mM lidocaine. **C)** SCC 4 cells were incubated with bitter agonists and XTT dye, an indicator of NADH production. A decrease in the difference of absorbance (475 nm –660 nm) indicates reduced NADH production. SCC 4 XTT absorbance values (475 nm – 660 nm) after 120 minutes of incubation with media, 1 – 10 mM lidocaine, or 1 -10 denatonium benzoate. Absorbance values were measured over six hours as seen in representative traces. **D)** SCC 47 XTT absorbance values (475 nm – 660 nm) after 6 hours of incubation with 10 mM lidocaine, 10 mM denatonium, or 1 μM – 100 μM ATP. Absorbance values were measured over six hours as seen in representative traces. Absorbance mean +/- SEM with >3 separate cultures. Significance by 1-way ANOVA with Bonferroni posttest comparing each treatment to media/control. $P < 0.05$ (*), $P < 0.01$ (**), $P < 0.001$ (***), and no statistical significance (ns or no indication).

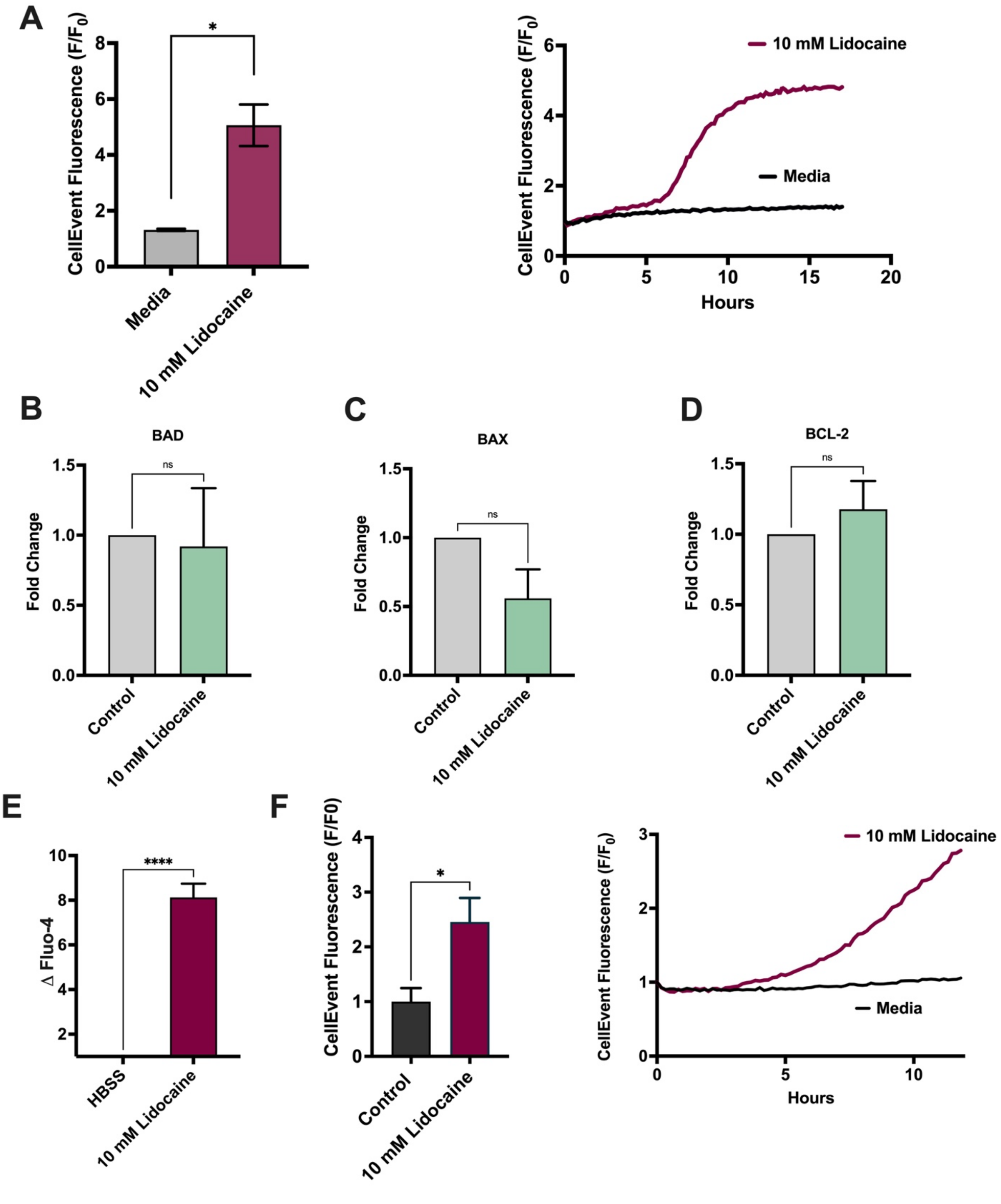


**Supplemental Figure 7. Lidocaine induces apoptosis in HNSCC and lung squamous cell carcinoma cells.** FaDu and SCC 47 cells were incubated with 10 mM lidocaine with CellEvent fluorescent dye, which fluoresces upon caspase-3 and caspase-7 cleavage. **A)** FaDu CellEvent fluorescence at 16 hours with 10 mM lidocaine. Representative trace of CellEvent fluorescence over 16 hours with 10 mM lidocaine. Fluorescent mean +/- SEM with >3 separate cultures. Significance determined by paired t-test between untreated and treated cells. **B-D)** SCC 47 cells were treated with 10 mM lidocaine for 6 hours. mRNA expression of *B)* BAD, *C)* BAX, and *D)* Bcl-2 were measured. 10 mM lidocaine normalized to untreated/control. Significance determined by paired t-test. **E)** NCI H520 (lung squamous cell carcinoma cell line) peak fluorescent $Ca^{2+}$ response with 10 mM lidocaine. Fluorescent mean +/- SEM with >3 separate cultures. Significance determined by unpaired t-test. **F)** NCI H520 CellEvent fluorescence at 16 hours with 10 mM lidocaine. Representative trace of CellEvent fluorescence over 16 hours with 10 mM lidocaine. Fluorescent mean +/- SEM with >3 separate cultures. Significance determined by paired t-test between untreated and treated cells. $P < 0.05$ (*), $P < 0.01$ (**), $P < 0.001$ (***), and no statistical significance (ns or no indication).

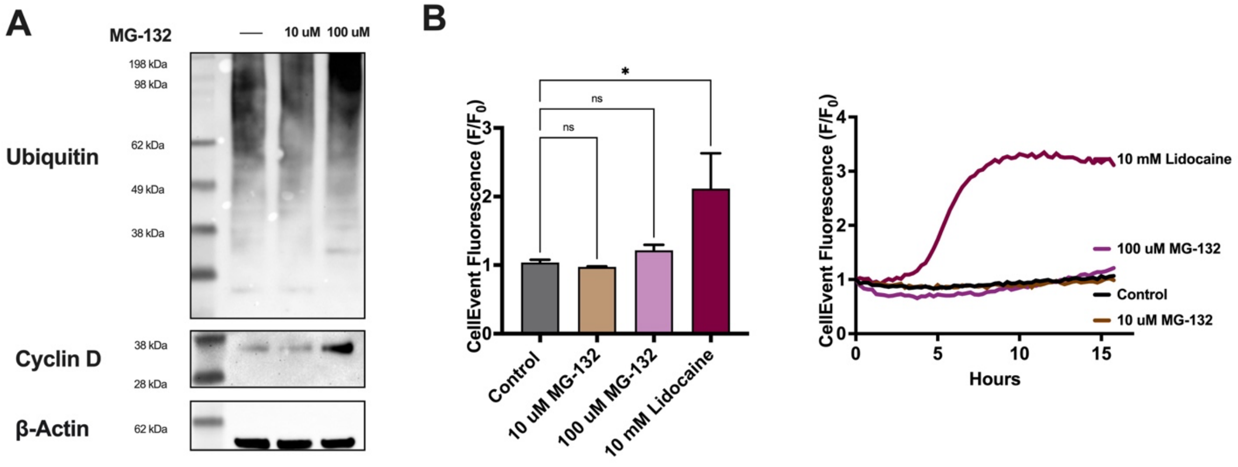


**Supplemental Figure 8. Proteasome inhibition with MG-132. A)** SCC 47 cells were treated with 10 or 100 uM MG-132 for 6 hours. Ubiquitin, β-Actin, and cyclin D primary antibody (1:1000) were used with anti-rabbit or anti-mouse IgG-horseradish peroxidase secondary antibodies (1:1000). **B)** SCC 47 cells were incubated with 0 – 100 uM MG-132 or 10 mM lidocaine with CellEvent fluorescent dye, which fluoresces upon caspase-3 and caspase-7 cleavage. Bare graph represents luorescent mean +/- SEM at 16 hours with >3 separate cultures. Trace represents CellEvent fluorescence over time. Significance determined by 1-way ANOVA comparing MG-132 and lidocaine treatment to control. P < 0.05 (*), P < 0.01 (**), P < 0.001 (***), and no statistical significance (ns or no indication).
